# A novel monomeric amyloid β-activated signaling pathway regulates brain development via inhibition of microglia

**DOI:** 10.1101/2024.06.13.598890

**Authors:** Hyo Jun Kwon, Devi Santhosh, Zhen Huang

## Abstract

Amyloid β (Aβ) forms aggregates in the Alzheimer’s disease brain and is well known for its pathological roles. Recent studies show that it also regulates neuronal physiology in the healthy brain. Whether Aβ also regulates glial physiology in the normal brain, however, has remained unclear. In this article, we describe the discovery of a novel signaling pathway activated by the monomeric form of Aβ in vitro that plays essential roles in the regulation of microglial activity and the assembly of neocortex during development in vivo. We find that activation of this pathway depends on the function of amyloid precursor (APP) and the heterotrimeric G protein regulator Ric8a in microglia and inhibits microglial immune activation at transcriptional and post-transcriptional levels. Genetic disruption of this pathway during neocortical development results in microglial dysregulation and excessive matrix proteinase activation, leading to basement membrane degradation, neuronal ectopia, and laminar disruption. These results uncover a previously unknown function of Aβ as a negative regulator of brain microglia and substantially elucidate the underlying molecular mechanisms. Considering the prominence of Aβ and neuroinflammation in the pathology of Alzheimer’s disease, they also highlight a potentially overlooked role of Aβ monomer depletion in the development of the disease.

## INTRODUCTION

Aβ, a core component of amyloid plaques in the Alzheimer’s disease brain, is well known to form oligomers under disease conditions. Studies have shown that the oligomers formed by Aβ are highly toxic, with wide-ranging effects including inhibition of neurotransmitter release, depletion of synaptic vesicle pools, disruption of postsynaptic organization and function, and impairment of multiple forms of synaptic plasticity (Gulisano et al., 2018; He et al., 2019; Kim et al., 2013; Lauren et al., 2009; Lazarevic et al., 2017; Parodi et al., 2010; Puzzo et al., 2008; Shankar et al., 2008; Walsh et al., 2002; Yang et al., 2015; Zott et al., 2019). These effects likely significantly underpin the pathogenic role of Aβ in Alzheimer’s disease and contribute to neuron loss and cognitive decline in patients. Besides its pathological roles, recent studies show that Aβ is also produced in the healthy brain by neurons in a neural activity-dependent manner and regulates the normal physiology of neurons (Cirrito et al., 2005; Fogel et al., 2014; Galanis et al., 2021; Garcia-Osta and Alberini, 2009; Gulisano et al., 2018; Gulisano et al., 2019; Morley et al., 2010; Palmeri et al., 2017; Puzzo et al., 2008; Zhou et al., 2022). For example, consistent with studies showing that Aβ monomers and low molecular weight oligomers positively regulate synaptic function and plasticity, administration of these molecules in vivo has been found to improve learning and memory in animals (Fogel et al., 2014; Garcia-Osta and Alberini, 2009; Gulisano et al., 2018; Gulisano et al., 2019; Morley et al., 2010; Palmeri et al., 2017; Puzzo et al., 2008). Furthermore, recent studies have shown that Aβ monomers directly promote synapse formation and function and homeostatic plasticity, processes crucial to normal cognitive function (Galanis et al., 2021; Kamenetz et al., 2003; Zhou et al., 2022). Together, these findings have provided crucial insights into the physiological roles that Aβ plays in regulating normal neuronal function in the brain. However, it remains unclear if Aβ also regulates the physiology of glia, nonneuronal cells that also play important roles in normal brain function.

Microglia and astrocytes, two of the major glial cell types in the brain, are known to play critical roles in the normal development, function, and plasticity of the brain circuitry (Barres, 2008; Schafer and Stevens, 2015). They coordinately regulate, among others, the spatiotemporally specific expression of immune cytokines in the brain that regulate numerous processes of brain circuit development, function, and plasticity (Zipp et al., 2023). For example, in the thalamus, a key relay station in the visual pathway, populations of astrocytes have been found to activate the expression of interleukin-33 in a neural activity-dependent manner, induce activity-dependent elimination of supernumerary synapses, and promote the maturation of the visual circuitry in early postnatal life (Vainchtein et al., 2018). In the adult hippocampus, in contrast, astrocytes have been found to activate the expression of interleukin-33 under neuronal activity blockade and induce homeostatic synaptic plasticity that maintains circuit activity balance (Wang et al., 2021). In the striatum and the neocortex, not only have astrocytes but also have microglia been observed to activate the expression of TNFα upon changes in neural circuit activity and induce homeostatic synaptic plasticity that dampens circuit perturbation (Heir et al., 2024; Lewitus et al., 2016; Stellwagen and Malenka, 2006). In the clinic, the induction of microglial release of cytokines such as TNFα also underpins the application of repetitive transcranial magnetic stimulation, a noninvasive brain stimulation technique frequently used to induce cortical plasticity and treat pharmaco-resistant depression (Eichler et al., 2023). In neurodegenerative diseases such as Alzheimer’s disease, glial activation and brain cytokine elevation are also key pathologic factors in disease development (Colonna and Butovsky, 2017; Patani et al., 2023). Furthermore, elevated TNFα expression by microglia also underlies interneuron deficits and autism-like phenotype linked to maternal immune activation (Yu et al., 2022). Thus, the precise regulation of glial cytokine expression in the brain plays a key role in the normal development and function of the brain and its dysregulation is linked to common neurodevelopmental and neurodegenerative diseases. However, how glial cytokine expression is mechanistically regulated by cell-cell communication in the brain have remained largely unknown.

In this article, we report the discovery of a novel microglial signaling pathway activated in vitro by Aβ, the neuron-produced peptide at the center of Alzheimer’s disease, that plays a crucial role in precisely regulating the levels of microglial cytokine expression and activity and ensuring the proper assembly of neuronal laminae during cerebral cortex development. We first came across evidence for this pathway in our study of the function of *ric8a*. *ric8a* encodes a guanine nucleotide exchange factor (GEF) and molecular chaperone for several classes of heterotrimeric G proteins, which become severely destabilized upon *ric8a* loss of function (Gabay et al., 2011; Papasergi-Scott et al., 2018; Tall et al., 2003). We found that deletion of *ric8a* during cortical development resulted in cortical basement membrane degradation, neuronal ectopia, and laminar disruption. However, unlike in classic models of cobblestone lissencephaly, these phenotypes resulted not from *ric8a* deficiency in brain neural cell types, but from deficiency in microglia. Ric8a-regulated Gα proteins are known to bind to the cytoplasmic domain of the amyloid precursor protein (APP) and mediate key branches of APP signaling in several cell types (Fogel et al., 2014; Milosch et al., 2014; Nishimoto et al., 1993; Ramaker et al., 2013). The *ric8a* cortical phenotypes also resemble those in triple or double mutants of APP family and pathway genes (Guenette et al., 2006; Herms et al., 2004), suggesting functional interactions. Indeed, we found that *app* deficiency in brain microglia also underpins ectopia formation in *app* family gene mutants. Furthermore, we found that APP and Ric8a form a pathway in microglia that is specifically activated by the monomeric form of Aβ and that this pathway normally inhibits the transcriptional and post-transcriptional expression of immune cytokines by microglia.

## RESULTS

### Cortical ectopia in *ric8a-emx1-cre* mutants results from non-neural deficiency

To study of the function of *ric8a,* a GEF as well as molecular chaperone for Gα proteins (Gabay et al., 2011; Papasergi-Scott et al., 2018; Tall et al., 2003), in neocortical development, we deleted a conditional *ric8a* allele (Ma et al., 2012; Ma et al., 2017) using *emx1-cre,* a *cre* line designed to target dorsal forebrain neural progenitors in mice (Gorski et al., 2002). We found it result in ectopia formation exclusively in the lateral cortex of the perinatal mutant brain (**Fig. 1a-d**). Birth-dating showed that the ectopia consisted of both early and late-born neurons (**Supplemental Fig. 1**). Consistent with this observation, neurons in the ectopia also stained positive for both Ctip2 and Cux1, genes specific to lower and upper-layer neurons, respectively. Interestingly, in cortical areas without ectopia, radial migration of early- and late-born neurons appeared largely normal as shown by birth-dating as well as Cux1 and Ctip2 staining (**Supplemental Fig. 2**). This suggests that cell-autonomous defects in neurons are unlikely the cause of the ectopia. At E16.5, clear breaches in the pial basement membrane of the developing cortex were already apparent (**Supplemental Fig. 3**). However, unlike classic models of cobblestone lissencephaly, where radial glial fibers typically retract, radial glial fibers in *ric8a* mutants instead extended beyond the breaches. This argues against radial glial cell adhesion defects since they would be predicted to retract. Furthermore, in areas without ectopia, we also observed normal localization of Cajal-Retzius cells, expression of Reelin, and splitting of the preplate, arguing against primary defects in Cajal-Retzius cells. In cobblestone lissencephaly, studies show that ectopia result from primary defects in radial glial maintenance of the pial basement membrane (Beggs et al., 2003; Graus-Porta et al., 2001; Moore et al., 2002; Satz et al., 2010). In *ric8a* mutants, we observed large numbers of basement membrane breaches at E14.5, almost all associated with ectopia (**Supplemental Fig. 4**). In contrast, at E13.5, although we also observed significant numbers of breaches, none was associated with ectopia. This indicates that basement membrane breaches similarly precede ectopia in *ric8a* mutants. However, at E12.5, despite a complete lack of basement membrane breaches, we observed increased numbers of laminin-positive debris across the lateral cortex, both beneath basement membrane segments with intact laminin staining and beneath segments with disrupted laminin staining, the latter presumably sites of future breach (**Supplemental Fig. 5**). As a major basement membrane component, the increased amounts of laminin debris suggest increased degradative activity within the developing cortex. Thus, these results indicate that excessive basement membrane degradation, but not defective maintenance, is likely a primary cause of cortical ectopia in *ric8a* mutants.

**Fig. 1.**
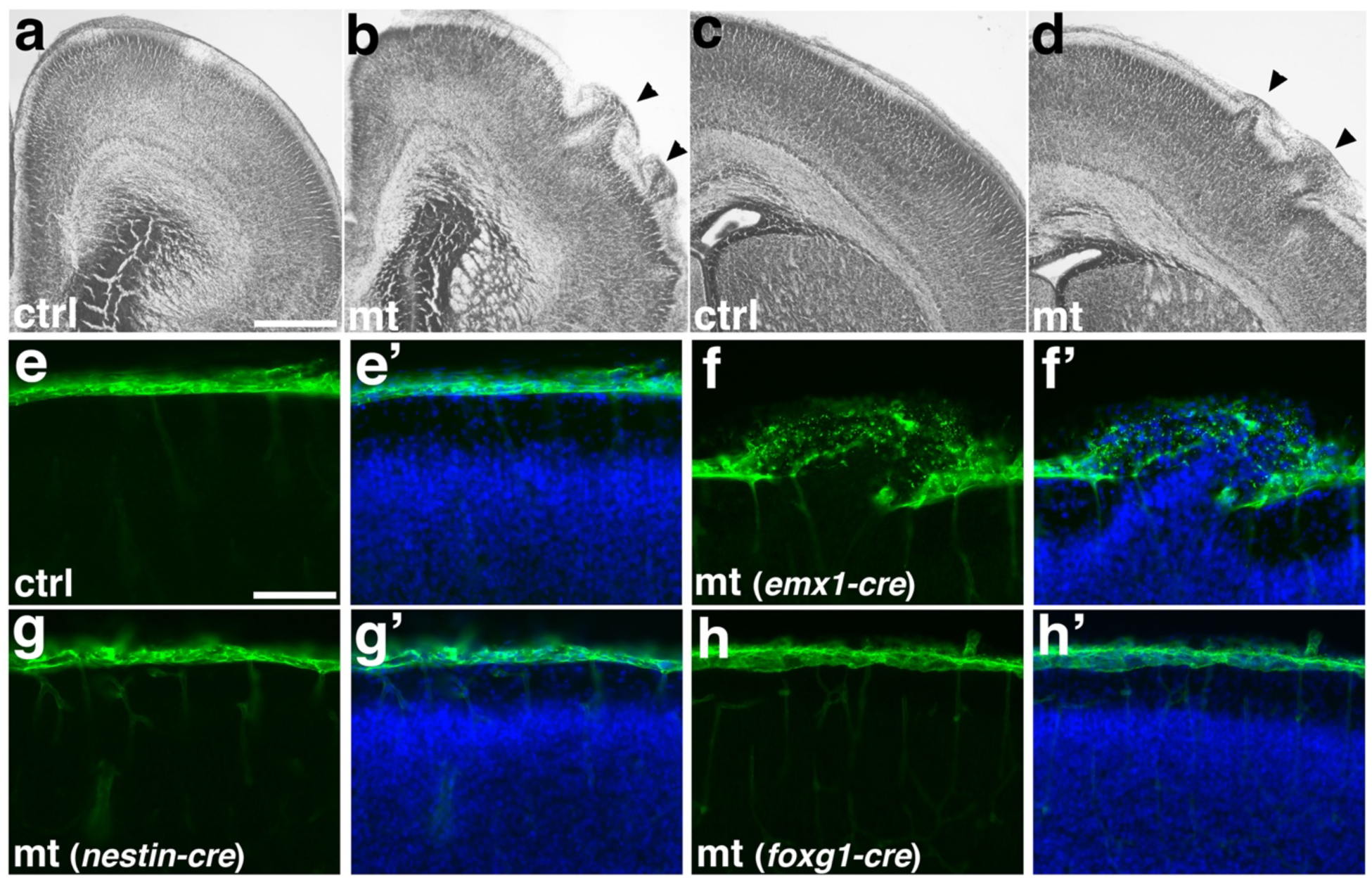
Deletion of *ric8a* using *emx1-cre* results in cortical ectopia due to non-neural deficits. (**a-d**) Nissl staining of control (ctrl, **a**&**c**) and mutant (mt, **b**&**d**) anterior motor (**a-b**) and posterior somatosensory (**c-d**) cortex at P0. **(e**-**e’**) Laminin (LN, in green) and nuclear (DAPI, in blue) staining of control cortices at P0. A continuous basement membrane is observed at the pia, beneath which cells are well organized in the cortical wall. (**f-f’**) Staining of *ric8a*-*emx1-cre* mutant cortices at P0. Basement membrane breach and neuronal ectopia are observed following *ric8a* deletion by *emx1-cre*, a *cre* line expressed in cortical radial glial progenitors beginning at E10.5. **(g**-**g’**) Staining of *ric8a*-*nestin-cre* mutant cortices at P0. No obvious basement membrane breach or neuronal ectopia is observed following *ric8a* deletion by *nestin-cre*, a *cre* line expressed in cortical progenitors beginning around E12.5. (**h-h’**) Staining of *ric8a*-*foxg1-cre* mutant cortices at P0. No obvious basement membrane breach or neuronal ectopia is observed following *ric8a* deletion by *foxg1-cre*, a *cre* line expressed in forebrain neural progenitors from E9.0. Scale bars, 640μm for (**a-b**), 400μm for (**c-d**), and 100μm for (**e-h’**).

To determine the cell type(s) genetically responsible for cortical basement membrane degradation and ectopia in *ric8a* mutants, we employed a panel of *cre* lines (**Fig. 1e-h’**). To target Cajal-Retzius cells, we employed *wnt3a-cre* (Yoshida et al., 2006) but found *ric8a* deletion using *wnt3a-cre* did not result in ectopia. To target postmitotic excitatory and inhibitory neurons, we employed *nex-cre* (Goebbels et al., 2006) and *dlx5/6-cre* (Stenman et al., 2003) respectively but similarly found neither result in ectopia. These results point to *ric8a* requirement in cell types other than post-mitotic neurons. To test the involvement of neural progenitors, we employed *nestin-cre* (Graus-Porta et al., 2001). Previous studies show that deletion of β*1 integrin* and related genes by *emx1-cre* and *nestin-cre* results in similar ectopia phenotypes (Belvindrah et al., 2006; Graus-Porta et al., 2001; Huang et al., 2006; Niewmierzycka et al., 2005). To our surprise, deletion of *ric8a* by *nestin-cre* did not result in ectopia (Ma et al., 2017) (**Fig. 1g-g’**). Since *nestin-cre*-mediated deletion in neural progenitors is inherited by post-mitotic neurons and astrocytes, this indicates that the combined deletion of *ric8a* from all these cell types does not lead to ectopia. The onset of *nestin-cre* expression is, however, developmentally slightly later than that of *emx1-cre* (Gorski et al., 2002). To assess the potential contribution of this temporal difference, we employed *foxg1-cre*, a *cre* line expressed in forebrain neural progenitors starting from E10.5 (Hebert and McConnell, 2000). We found that *ric8a* deletion using *foxg1-cre* still failed to produce ectopia (**Fig. 1h-h’**). Thus, these results strongly argue against the interpretation that *ric8a* deficiency in neural cell lineages is responsible for basement membrane degradation and ectopia in *ric8a* mutants.

During embryogenesis, the neural tube undergoes epithelial-mesenchymal transition giving rise to neural crest cells (Leathers and Rogers, 2022). This process involves region-specific basement membrane breakdown that resembles the *ric8a* mutant phenotype. To determine if ectopic epithelial-mesenchymal transition plays a role, we examined potential changes in neuro-epithelial cell fates in the mutant cortex. We found that cortical neural progenitors expressed Pax6, Nestin, and Vimentin normally (**Supplemental Fig. 6**). Cell proliferation in the ventricular zone was also normal. Furthermore, although *ric8a* regulates asymmetric cell division in invertebrates (Afshar et al., 2004; Couwenbergs et al., 2004; David et al., 2005; Hampoelz et al., 2005; Wang et al., 2005), we observed no significant defects in mitotic spindle orientation at the ventricular surface. Additionally, no ectopic expression of neural crest markers or Wnt pathway activation was observed (**Supplemental Fig. 7**). Altogether, these results further indicate that non-neural-cell deficiency is responsible for ectopia formation in *ric8a* mutants.

### Microglial *ric8a* deficiency is responsible for ectopia formation

To assess the role of non-neural cell types, we turned our attention to microglia since RNA-seq studies show that brain microglia express *emx1* at a significant level (Zhang et al., 2014). To determine if *emx1-cre* is expressed and active in microglia, we isolated microglia from *ric8a-emx1-cre* mutants. We found that *emx1-cre* mediated *ric8a* deletion resulted in altered cytokine expression in microglia (**Supplemental Fig. 8a-b**). This indicate that *emx1-cre* is expressed and active in microglia and deletes *ric8a*. In further support of this interpretation, we found that when crossed to a reporter, *emx1-cre* resulted in the expression of reporter gene in microglia (**Supplemental Fig. 8c-c”**). It also resulted in the reduction of *ric8a* mRNA levels in in microglia in *ric8a-emx1-cre* mutants (**Supplemental Fig. 8d**). To determine the specific significance of *ric8a* deletion from microglia alone, we next employed a microglia-specific *cx3cr1-cre* (Yona et al., 2013). Like *emx1-cre* mutants, *ric8a-cx3cr1-cre* mutant microglia also showed elevated cytokine secretion and transcription in comparison to control microglia upon stimulation by lipopolysaccharide (LPS) (**Fig. 2a-b**). Similar results were also obtained with stimulation by polyinosinic-polycytidylic acid (poly I:C), an intracellular immune activator. Thus, these results indicate that *ric8a* deficiency in microglia results in broad increases in microglial sensitivity to immune stimulation.

**Fig. 2.**
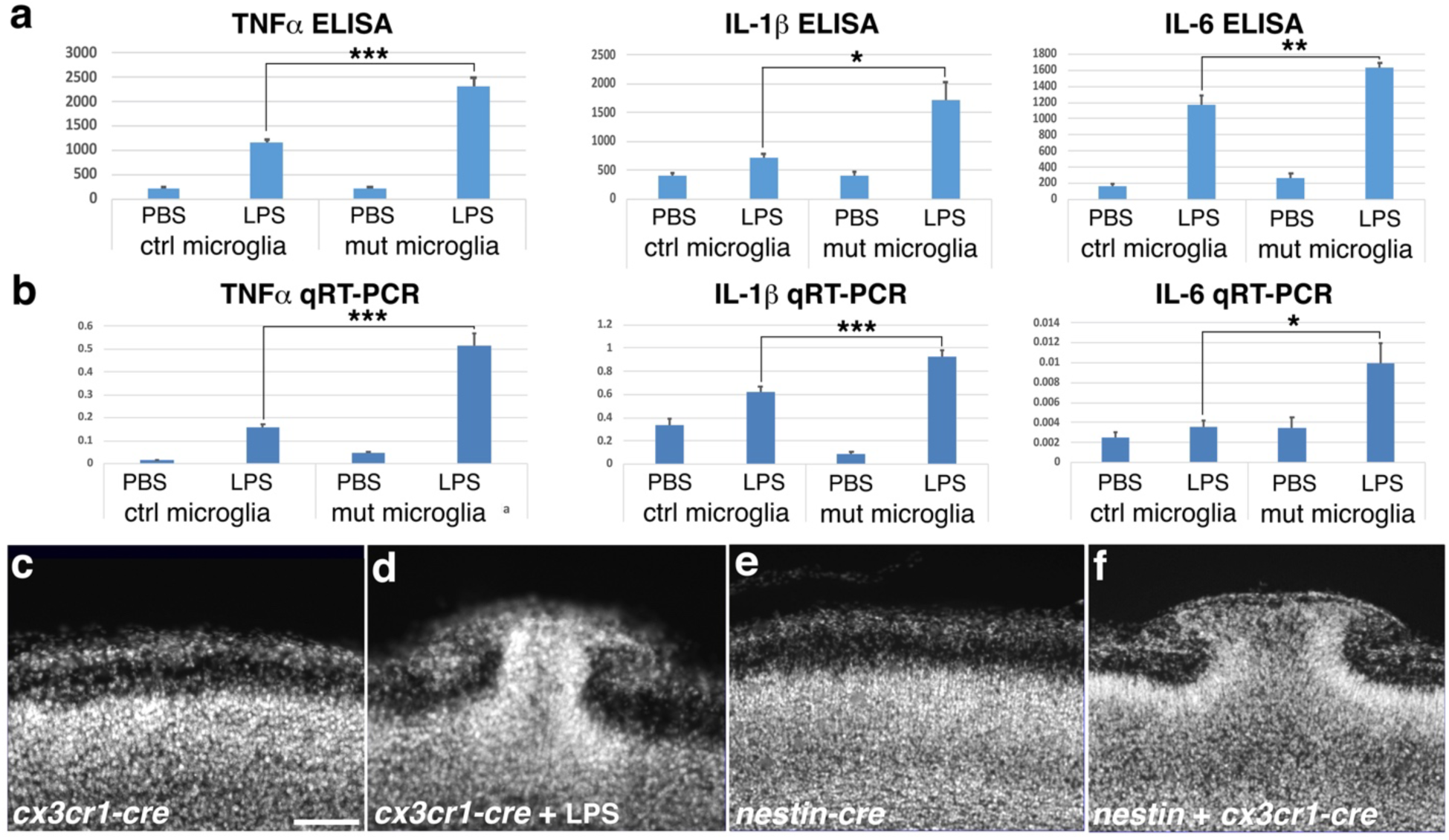
*ric8a* deficiency in microglia is responsible for cortical ectopia. **(a)** TNFα, IL-1β, and IL-6 secretion (pg/ml) in control and *ric8a/cx3cr1-cre* mutant microglia following LPS stimulation. *, *P* < 0.05; **, *P* < 0.01; ***, *P* < 0.001; n = 6-8 each group. **(b)** TNFα, IL-1 β, and IL-6 mRNA expression in control and *ric8a/cx3cr1-cre* mutant microglia following LPS stimulation. *, *P* < 0.05; ***, *P* < 0.001; n = 5-6 each group. (**c**-**d**) Nuclear (DAPI, in grey) staining of *ric8a-cx3cr1-cre* mutant cortices at P0 in the absence (**c**) or presence (**d**) of LPS treatment during embryogenesis. (**e-f**) Nuclear (DAPI, in grey) staining of *ric8a-nestin-cre* single *cre* (**e**) and *ric8a-nestin-cre+cx3cr1-cre* double *cre* (**f**) mutant cortices at P0. Scale bar in (**c**), 100 μm for (**c**-**f**).

To determine if microglial *ric8a* deficiency alone is sufficient to cause cortical ectopia in vivo, we examined *ric8a-cx3cr1-cre* mutants but found that it did not affect either basement membrane integrity or cortical layering (**Fig. 2c**). We reasoned that this may be related to the fact that *ric8a* mutant microglia only show heightened activity upon stimulation but not under basal unstimulated conditions (**Fig. 2a-b**) but elevated microglial activity may be needed for basement membrane degradation and ectopia formation. To test this possibility, we employed in utero LPS administration to activate microglia during cortical development. We found that over 50% of *ric8a-cx3cr1-cre* mutant neonates showed ectopia when administered LPS at E11.5-12.5 (10 of 19 mutant neonates examined) (**Fig. 2d**). In contrast, no cortical ectopia were observed in any of the 32 littermate controls that were similarly administered LPS. This indicates that only the combination of microglial *ric8a* deficiency and immune activation leads to ectopia formation. In *emx1-cre* mutants, ectopia develop without LPS administration (**Fig. 1**). We suspect that this may be due to concurrent *ric8a* deficiency in neural cell types, which may result in deficits that mimic immune stimulation. In the embryonic cortex, for example, studies have shown that large numbers of cells die starting as early as E12 (Blaschke et al., 1996; Blaschke et al., 1998). Radial glia and neuronal progenitors play critical roles in the clearance of apoptotic cells and cellular debris in the brain (Amaya et al., 2015; Ginisty et al., 2015; Lu et al., 2011) and Ric8a-dependent heterotrimeric G proteins promotes this function in both professional and non-professional phagocytic cells (Billings et al., 2016; Flak et al., 2020; Pan et al., 2016; Preissler et al., 2015; Zhang et al., 2023). Thus, *ric8a* deficiency in radial glia may potentially result in accumulation of apoptotic cell debris in the embryonic brain that stimulate microglia. To test this, we next additionally deleted *ric8a* from radial glia in the *ric8a*-*cx3cr1-cre* microglial mutant background by introducing *nestin-cre*. We have shown that *ric8a* deletion by *nestin-cre* alone does not result in ectopia (**Fig. 1g-g’**). However, we found that, like deletion by *emx1-cre*, *ric8a* deletion by the dual *cre* combination of *cx3cr1-cre* and *nestin-cre* also resulted in severe ectopia in all double *cre* mutants (6 of 6 examined) (**Fig. 2f**). Thus, these results indicate that elevated immune activation of *ric8a* deficient microglia during cortical development is responsible for ectopia formation.

### Microglial *app* deficiency also results in ectopia formation

In the large numbers of cobblestone lissencephaly and related cortical ectopia mutants, besides the lateral cortex, severe ectopia are typically also observed at the cortical midline (Beggs et al., 2003; Belvindrah et al., 2006; Graus-Porta et al., 2001; Huang et al., 2006; Moore et al., 2002; Niewmierzycka et al., 2005; Satz et al., 2010). There are only a few mutants including the *ric8a-emx1cre* mutant that are exception, in that the ectopia are not observed at the cortical midline but are instead exclusively located to the lateral cortex (**Fig. 1**). The other mutants in this unique group include the *app/aplp1/2* triple (Herms et al., 2004) and *apbb1/2* double knockouts (Guenette et al., 2006). This suggests that similar mechanisms involving microglia may underlie ectopia formation in *ric8a-emx1cre, app/aplp1/2,* and *apbb1/2* mutants. Independent studies also point to a role of non-neuronal cells in ectopia formation in *app* family gene mutants. For example, unlike the triple knockout, which causes neuronal over-migration, specific *app* knockdown in cortical neurons during development results in under-instead of over-migration of targeted neurons (Young-Pearse et al., 2007). Furthermore, Ric8a-regulated Gα proteins play a conserved role in mediating key branches of APP signaling in cells across species (Fogel et al., 2014; Milosch et al., 2014; Nishimoto et al., 1993; Ramaker et al., 2013) and we confirmed that Gαi proteins are severely depleted in *ric8a-emx1-cre* mutant cortices (**Supplemental Fig. 9**). Thus, like in *ric8a-emx1cre* mutants, microglia may play a key role in ecotopia formation in APP pathway mutants. To test this, we first analyzed *app* mutant microglia. To this end, we employed *cx3cr1-cre* to delete a conditional allele of *app* from microglia and found that microglia cultured from *app*-*cx3cr1-cre* mutants showed reduced TNFα and IL-6 secretion as well as muted IL-6 transcription upon stimulation (**Fig. 3a, Supplemental Fig. 10a-b**). This indicates that *app* plays a previously unrecognized, cell-autonomous role in microglia in regulating microglial activity. Microglia exhibit attenuated immune response following chronic stimulation, especially when carrying strong loss-of-function mutations in anti-inflammatory pathways (Chamberlain et al., 2015; Sayed et al., 2018). We suspect that the attenuated response by *app* mutant microglia may result from similar effects following in vitro culture. To test effects of *app* mutation under conditions that more closely resemble in vivo physiological conditions, we next isolated fresh, unelicited peritoneal macrophages and acutely analyzed their response to immune stimulation. We found that *app* mutant macrophages showed significantly elevated secretion of all cytokines tested (**Fig. 3b**). At the transcriptional level, mRNA induction was also increased for all cytokines (**Fig. 3c**). Thus, like that of *ric8a*, the normal function of *app* also appears to be to suppress the inflammatory activation of microglia.

**Fig. 3.**
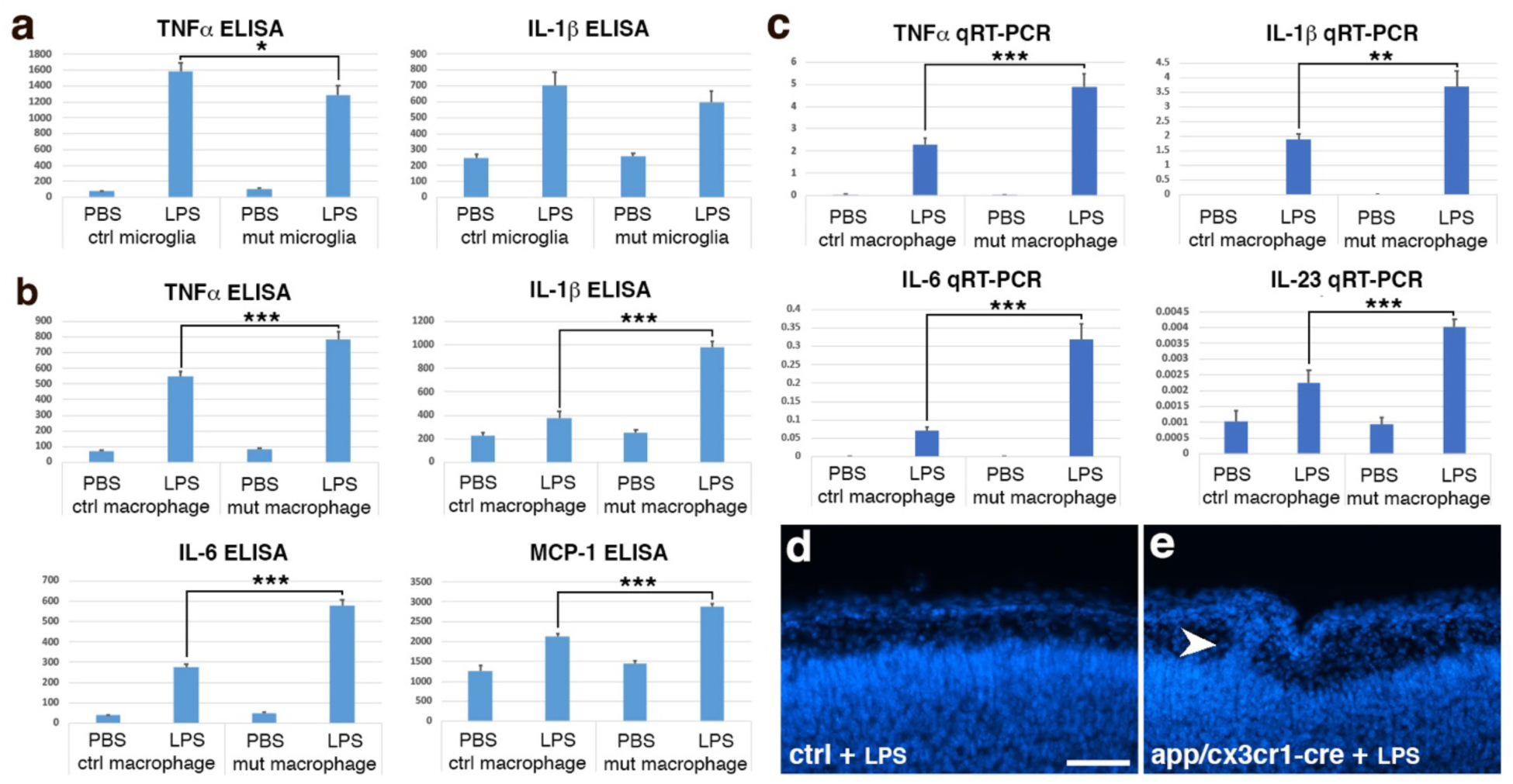
*app* deficiency results in hypersensitive microglia and cortical ectopia. **(a)** TNFα and IL-1β secretion (pg/ml) in cultured control and *app/cx3cr1-cre* mutant microglia following LPS stimulation. *, *P* < 0.05; n = 7-9 each group. **(b)** TNFα, IL-1β, IL-6, and MCP1 secretion (pg/ml) in fresh unelicited control and *app/cx3cr1-cre* mutant peritoneal macrophages following LPS stimulation. ***, *P* < 0.001; n = 7-10 each group. **(c)** TNFα, IL-1 β, IL-6, and IL-23 mRNA expression in fresh unelicited control and *app/cx3cr1-cre* mutant peritoneal macrophages following LPS stimulation. **, *P* < 0.01; ***, *P* < 0.001; n = 6 each group (**d-e**) Nuclear (DAPI, in blue) staining of control (**D**) and LPS-treated *app/cx3cr1-cre* mutant (**E**) cortices at P0. Note cortical ectopia in the mutant cortex (arrowhead). Scale bar in (**d**), 200μm for (**d-e**).

To determine if microglial *app* deficiency is also responsible for ectopia formation in *app* triple knockout mutants, we next asked if activating microglia in microglia-specific *app* mutants similarly results in pial ectopia during cortical development. To this end, we administered LPS in utero at E11.5-12.5 to *app-cx3cr1-cre* mutant animals as we did to *ric8a-cx3cr1-cre* mutants above. We found that, while none of the 81 littermate controls administered LPS showed ectopia, a significant number of mutant neonates showed ectopia (6 of 31 neonates examined, ∼19%) and associated breaches in the basement membrane (**Fig. 3e, Supplemental Fig. 10c-f”**). Thus, *app* deficient microglia, when activated, also results in cortical ectopia during development. The reduced severity of the ectopia observed, as compared to that in *ric8a/cx3cr1-cre* mutants, likely in part results from the reduced LPS dosage (by ∼3 folds) we had to use in these animas due to the enhanced immune sensitivity of their strain genetic background. Other *app* gene family members are also expressed in microglia (Zhang et al., 2014) and ectopia are only observed in *app/aplp1/2* triple but not in any double or single mutants (Herms et al., 2004). Aplp1/2 may therefore also compensate for the loss of APP in microglia. Thus, these results indicate that *app* normally plays a cell-autonomous role in microglia that negatively regulate microglial activation, and its loss of function underlies cortical ectopia formation. The similarities between *app* and *ric8a* mutant phenotypes suggest that they form a previously unknown anti-inflammatory pathway in microglia.

### Monomeric Aβ suppresses microglial inflammatory activation via an APP-Ric8a pathway

The possibility that *app* and *ric8a* may form a novel anti-inflammatory pathway in microglia raises questions on the identity of the ligands for the pathway. Several molecules have been reported to bind to APP and/or activate APP-dependent pathways (Fogel et al., 2014; Milosch et al., 2014; Rice et al., 2012), among which Aβ is noteworthy for its nanomolar direct binding affinity (Fogel et al., 2014; Shaked et al., 2006). Aβ oligomers and fibrils have been shown by numerous studies to be pro-inflammatory, while non-fibrillar Aβ lack such activity (Halle et al., 2008; Huang, 2023, 2024; Lorton et al., 1996; Muehlhauser et al., 2001; Tan et al., 1999). In contrast, when employed under conditions that favor the monomer conformation, Aβ inhibits T cell activation (Grant et al., 2012). This suggests that, unlike pro-inflammatory Aβ oligomers (**Supplemental Fig. 11j**), Aβ monomers may be anti-inflammatory. To test this possibility, we dissolved Aβ40 peptides in DMSO, a standard approach in Alzheimer’s disease research that has been shown to preserve the monomeric conformation (LeVine, 2004; Stine et al., 2011). We found that Aβ monomers as prepared potently suppressed the secretion of large numbers of cytokines (**Fig. 4a**, **Supplemental Fig. 10**) and showed similar effects on microglia no matter if they were activated by LPS or poly I:C (**Fig. 4b**). We also found that the Aβ monomers similarly strongly inhibited the induction of cytokines at the transcriptional level (**Fig. 4c, Supplemental Fig. 11**). In addition, we observed these effects with Aβ40 peptides from different commercial sources. Thus, these results indicate that monomeric Aβ possesses a previously unreported anti-inflammatory activity against microglia that strongly inhibits microglial inflammatory activation.

**Fig. 4.**
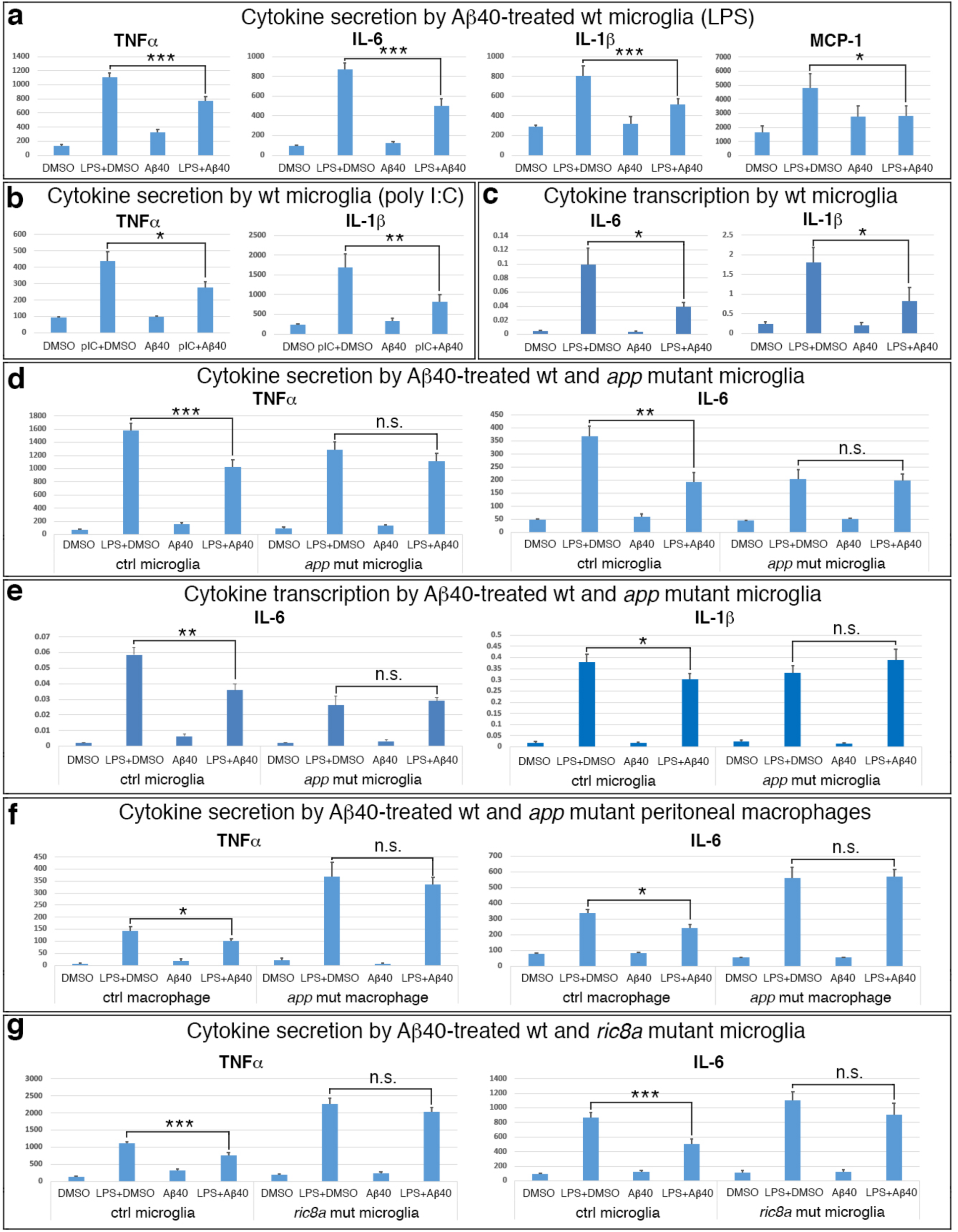
Monomeric Aβ40 suppresses microglia via an APP and Ric8a. (a) TNFα, IL-6, IL-1β, and MCP1 secretion (pg/ml) by wildtype microglia following LPS stimulation in the absence or presence of Aβ40 (200 or 500nM). *, *P* < 0.05; ***, *P* < 0.001; n = 8-14 each group. (b) TNFα and IL-1β secretion (pg/ml) by wildtype microglia following poly I:C stimulation in the absence or presence of Aβ40 (500nM). *, *P* < 0.05; **, *P* < 0.01; n = 6-7 each group. (c) IL-6 and IL-1β mRNA induction in wildtype microglia following LPS stimulation in the absence or presence of Aβ40 (500nM). *, *P* < 0.05; n = 6 each group. (d) TNFα and IL-6 secretion (pg/ml) by control and *app/cx3cr1-cre* mutant microglia following LPS stimulation in the absence or presence of Aβ40 (200nM). **, *P* < 0.01; ***, *P* < 0.001; n = 8 each group. (e) IL-6 and IL-1β mRNA induction in control and *app/cx3cr1-cre* mutant microglia following LPS stimulation in the absence or presence of Aβ40 (200nM). *, *P* < 0.05; **, *P* < 0.01; n = 6 each group. (f) TNFα and IL-6 secretion (pg/ml) by control and *app/cx3cr1-cre* mutant peritoneal macrophages following LPS stimulation in the absence or presence of Aβ40 (500nM). *, *P* < 0.05; n = 6-7 each group (g) TNFα and IL-6 secretion (pg/ml) by control and *ric8a/cx3cr1-cre* mutant microglia following LPS stimulation in the absence or presence of Aβ40 (200nM). ***, *P* < 0.001; n = 12-14 each group.

To determine whether monomeric Aβ signals through APP, we employed *app-cx3cr1-cre* mutant microglia. We found that, unlike that of control microglia, Aβ monomers failed to suppress the secretion of all tested cytokines by *app* mutant microglia (**Fig. 4d, Supplemental Fig. 11**). Interestingly, this blockade appeared to be specific to *app* since Aβ monomers still significantly suppressed cytokine secretion by *aplp2* mutant microglia. At the transcriptional level, Aβ monomers also failed to suppress cytokine induction in *app* mutant microglia (**Fig. 4e, Supplemental Fig. 11**). Together, these results indicate that APP is functionally required in microglia for Aβ monomer inhibition of cytokine expression at both transcriptional and post-transcriptional levels. Cultured microglia from *app-cx3cr1-*cre mutants showed attenuated immune activation (**Fig. 3**). To assess whether this may affect the efficacy of Aβ monomer inhibition, we next tested the response of fresh, unelicited macrophages. We found that, like that of control microglia, cytokine secretion by control macrophages was also strongly suppressed by Aβ monomers (**Fig. 4f, Supplemental Fig. 11**). However, even though *app* mutant macrophages showed elevated response to immune stimulation in comparison to control macrophages, they still failed to respond to Aβ monomers and displayed levels of cytokine secretion that were indistinguishable from those of DMSO-treated cells (**Fig. 4f, Supplemental Fig. 11**). Thus, these results further indicate that APP function is required in microglia for mediating the anti-inflammatory effects of Aβ monomers.

The similarity of *ric8a* ectopia to *app* ectopia phenotype (**Figs. 2** & **3**) also suggests that Ric8a functions in the same pathway as APP in mediating Aβ monomer anti-inflammatory signaling in microglia. This is consistent with previous studies showing that heterotrimeric G proteins are coupled to APP and mediate APP intracellular signaling in vitro and vivo (Fogel et al., 2014; Milosch et al., 2014; Nishimoto et al., 1993; Ramaker et al., 2013) and that Ric8a is a molecular chaperone essential for the post-translational stability of heterotrimeric G proteins (Gabay et al., 2011; Tall et al., 2003). To directly test if Ric8a is part of this pathway, we next employed *ric8a-cx3cr1-cre* mutant microglia. We found that, indeed, like that of *app* mutant microglia, Aβ monomers also failed to suppress the secretion of TNFα and IL-6 by *ric8a* mutant microglia (**Fig. 4g**). This indicates that heterotrimeric G proteins function is likely required in the same pathway of APP in microglia for the suppression of TNFα and IL-6 secretion. However, unlike APP, we found that Ric8a appears to be dispensable for Aβ monomer regulation of other cytokines. For example, unlike that of TNFα and IL-6, Aβ monomers still suppressed IL-1β secretion by *ric8a* mutant microglia (**Supplemental Fig. 11**). It also appears to be dispensable for the regulation of cytokine transcription since Aβ monomers similarly suppressed IL-6 transcriptional induction in both control and *ric8a* mutant microglia. These results suggest that heterotrimeric G proteins function may only mediate some of the anti-inflammatory signaling of monomeric Aβ. Thus, APP and Ric8a-regulated heterotrimeric G proteins form part of a novel anti-inflammatory pathway activated by monomeric Aβ in microglia.

### Elevated matrix metalloproteinases (MMPs) cause basement membrane degradation

We have shown that heightened microglial activation due to mutation in the Aβ monomer-activated APP/Ric8a pathway results in basement membrane degradation and ectopia during cortical development. To further test this interpretation, we sought to test the prediction that inhibition of microglial activation in these mutants suppressed the formation ectopia. To this end, we employed dorsomorphin and S3I-201, inhibitors targeting Akt, Stat3, and other mediators in pro-inflammatory signaling (Lee et al., 2016; Qin et al., 2012). Consistent with their anti-inflammatory activity, we found that dorsomorphin and S3I-201 both suppressed astrogliosis associated with neuroinflammation in the cortex of *ric8a-emx1*-*cre* mutants (**Supplemental Fig. 12**). Furthermore, they also suppressed the formation of ectopia in *ric8a-emx1*-*cre* mutants, reducing both the number and the size of the ectopia observed (**Fig. 5a-f**). Most strikingly, the combined administration of dorsomorphin and S3I-201 nearly eliminated all ectopia in *ric8a-emx1*-*cre* mutants (**Fig. 5d, 5e**). Thus, these results indicate that excessive inflammatory activation of microglia is responsible for ectopia formation in *ric8a* mutants.

**Fig. 5.**
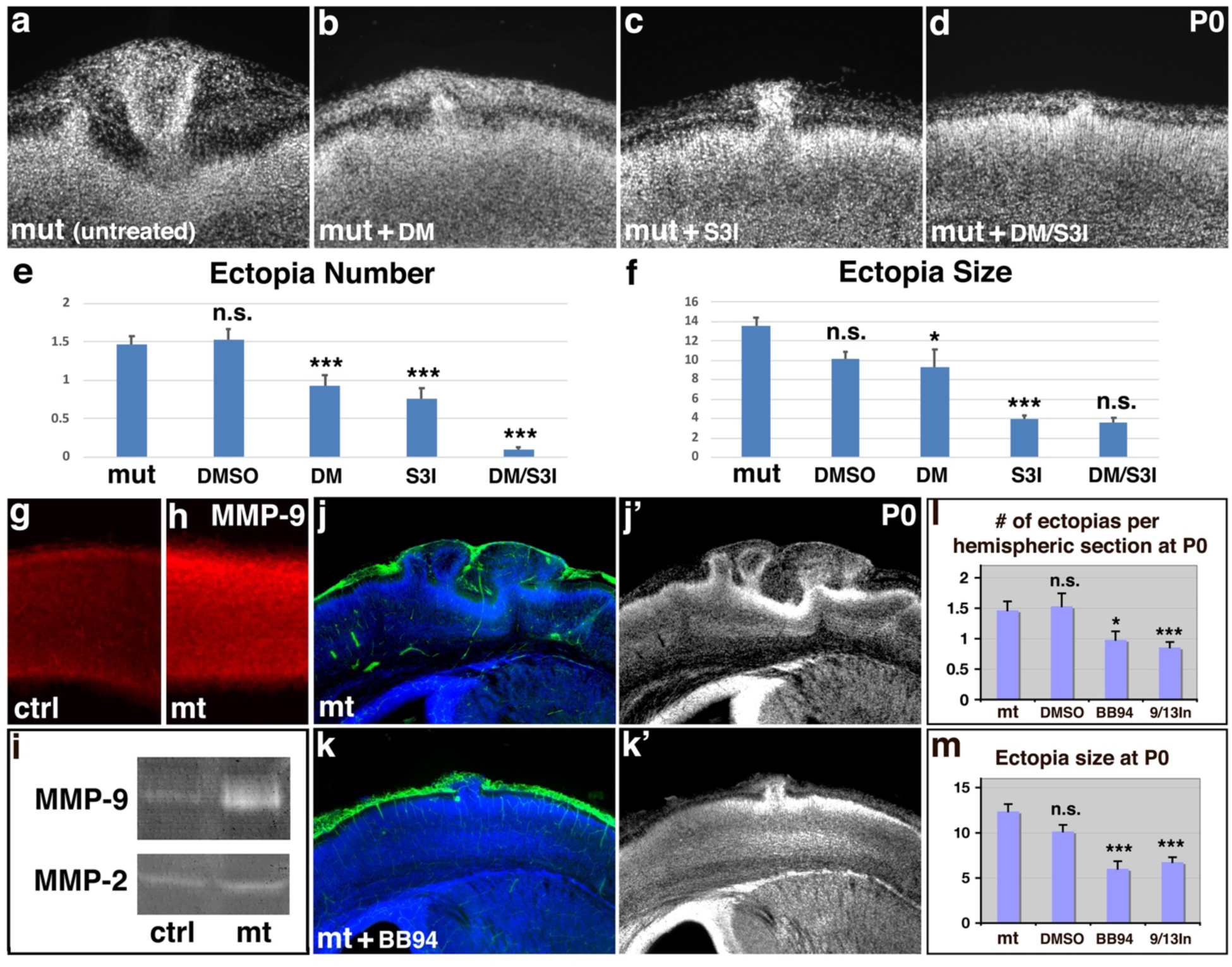
Inhibition of both microglial inflammatory activation and cortical MMP9 activity suppresses basement membrane breach and neuronal ectopia. **(a-d**) Nuclear (DAPI, in grey) staining of untreated (**a**), anti-inflammatory drug dorsomorphin (DM) (**b**), Stat3 inhibitor S3I-201 (S3I) (**c**), and DM/S3I (**d**) dual treated mutant cortices at P0. **(e-f**) Quantitative analysis of ectopia number (**e**) and size (**f**) in the neonatal mutant cortex after DMSO, DM, S3I, and DM/S3I dual treatment at E12.5. *, *P* < 0.05; ***, *P* < 0.001; all compared to untreated mutants. The reduction in ectopia size after dual treatment is not statistically significant, likely due to the small number of ectopias that remained. (**g**-**h**) MMP9 (in red) staining of control (**g**) and mutant cortices (**h**) at E13.5. Quantification shows statistically significant increases in mutants (control, 24.8 ± 0.2 AU (Arbitrary Units); mutant, 35.7 ± 1.7 AU; *P* = 0.002; n = 6). (**i**) Gel zymography of control and mutant cortical lysates at E13.5. Increased levels of MMP9 but not of MMP2 were observed in mutants (control, 1.00 ± 0.06 AU; mutant, 3.72 ±1.86 AU; *P* = 0.028; n = 4). See further details in supplemental Fig. 13. (**j-k’**) Laminin (in green) and nuclear (DAPI, in blue) staining of mutant cortices untreated (**H**) or treated (**I**) with BB94. (**l-m**) Quantitative analysis of ectopia number and size following MMP inhibitor BB94 or MMP9/13 inhibitor I treatment. *, *P* < 0.05; ***, *P* < 0.001; all compared to untreated mutants.

Under neuroinflammatory conditions, brain cytokines frequently induce MMPs, which lead to breakdown of the extracellular matrix and contribute to disease pathology (Pagenstecher et al., 1998; Wang et al., 2000). Since *ric8a* mutant microglia are hyperactive in inflammatory cytokine production, we wonder if induction of MMPs may underlie the laminin degradation and cortical basement membrane break observed in *ric8a* mutants. To test this, we examined MMP9 expression in the embryonic cortex by in situ hybridization. We found that at E13.5, MMP9 mRNA is strongly expressed in a sparse cell population resembling microglia populating the cortex at this stage (Squarzoni et al., 2014) (**Supplemental Fig. 13**). Next, we examined the activities of MMP2 and MMP9 in the developing control and mutant cortex using gelatin gel zymography. We found that the activity of MMP9 in the mutant cortex was significantly increased (**Fig. 5i, Supplemental Fig. 13**). In contrast, that activity of MMP2 remained unaffected. Similarly, at the protein level, we found that the immunoreactivity for MMP9 was increased in *ric8a-emx1*-*cre* mutants (**Fig. 5g-h**). To test if the increased MMP activity is responsible for the ectopia observed, we next employed BB94, a broad-spectrum inhibitor of MMPs. We found that BB94 administration significantly suppress both the number and the size of the ectopia in *ric8a* mutants (**Fig. 5j-m**). To narrow down the identity of MMPs responsible, we further employed an inhibitor specific for MMP9 and 13 (MMP-9/MMP-13 inhibitor I, CAS 204140-01-2) and found that it similarly suppressed both the number and the size of the ectopia (**Fig. 5l-m**). Furthermore, consistent with its near complete suppression of cortical ectopia (**Fig. 5a-f**), we found that the co-administration of dorsomorphin and S3I-201 also reduced MMP9 activity in the mutant cortex to the control level (**Supplemental Fig. 14**). Thus, these results indicate this Aβ monomer-regulated anti-inflammatory pathway normally promotes cortical development through suppressing microglial activation and MMP induction.

## DISCUSSION

The spatiotemporal expression of immune cytokines by glial cells in the brain plays critical roles in the normal development, function, and plasticity of the brain circuitry (Barres, 2008; Schafer and Stevens, 2015; Zipp et al., 2023). In this article, we have identified a novel microglial anti-inflammatory pathway activated by monomeric Aβ that inhibits microglial cytokine expression and plays essential roles in the normal development of the cerebral cortex. We have found that this pathway is mediated by APP and the heterotrimeric G protein GEF and molecular chaperone Ric8a in microglia and its activation leads to the inhibition of microglial cytokine induction at transcriptional and post-transcriptional levels (**Figs. 1-4**). We further show that a key function of this pathway is to suppress the activity of MMP9 during corticogenesis and disruption of this regulation results in cortical basement membrane degradation and neuronal ectopia development (**Figs. 1-3, 5**). Furthermore, we find that this pathway is activated specifically by the monomeric form of Aβ in vitro (**Fig. 4**), identifying, for the first time, an isoform specific activity of Aβ against microglia. These results provide novel insights into the neuron-glia communication mechanisms that coordinate the regulation of immune cytokines, key regulators of Hebbian and non-Hebbian synaptic plasticity, by glial cells in the brain. The discovery of the novel activity of monomeric Aβ as a negative regulator of microglia may also facilitate the further elucidation of Alzheimer’s disease pathogenesis.

### Microglial activity regulation during cortical development

Among the glial cell populations in the brain, astrocytes and oligodendrocyte are both born within the nervous system at the end of cortical neurogenesis. As such, they play limited roles in the early steps of cortical development. In contrast, microglia are not only of a distinct non-neural lineage that originates from outside the nervous system but also begin to populate the brain at the onset of corticogenesis (Ginhoux et al., 2010; Hattori et al., 2023). As such, they play unique roles throughout cortical development. Indeed, microglial activity has been found to regulate the size of the cortical neural precursor pool (Cunningham et al., 2013). Microglia-secreted cytokines have also been found to promote both neurogenesis and oligodendrogenesis (Shigemoto-Mogami et al., 2014). As such, the precise regulation of microglial activity is critical to the normal development of the neocortex from an early stage. In this study, we have shown that immune over-activation of microglia deficient in a monomeric Aβ-regulated pathway results in excessive cortical matrix proteinase activation, leading basement membrane degradation and neuronal ectopia. Previous studies have shown that reductions in the expression of microglial immune and chemotaxis genes instead lead to the failure of microglia to populate the brain (Iyer et al., 2022). These results together highlight the importance of precisely regulating the level of microglial activity during brain development. The dramatic destructive effects of microglial hyperactivity observed during corticogenesis also foreshadow the critical roles it plays in brain dysfunction and disease at later stages of life.

In this study, we have also shown that the anti-inflammatory regulation of microglia in corticogenesis depends on a pathway composed of APP and the heterotrimeric G protein regulator Ric8a. This has revealed new insight into the intercellular signaling mechanisms regulating microglial activity in the brain. Heterotrimeric G proteins are well-known mediators of G protein-coupled receptor (GPCR) signaling. In this study, we have found that they likely also function in the same pathway as APP. To our knowledge, ours is the first study to report an in vivo anti-inflammatory function of this pathway in microglia and has significantly advanced knowledge in microglial biology. This is also consistent with previous studies showing that heterotrimeric G proteins directly interact with the APP cytoplasmic domain and mediate key branches of APP signaling from invertebrates to mammals (Fogel et al., 2014; Milosch et al., 2014; Nishimoto et al., 1993; Ramaker et al., 2013). In this study, we have in addition shown that this pathway is specifically activated in vitro by the monomeric form of Aβ, a peptide produced by neurons in the brain (Cirrito et al., 2005), providing further insight into the biological function of this pathway. In the early cortex, neurogenesis is just beginning, and most neurons born are in an immature state. It is unclear if this pathway is activated by Aβ at this stage in vivo. However, studies have shown that other APP ligands such as pancortin, a member of the olfactomedin family proteins known to inhibit innate immunity (Liu et al., 2010), are expressed in the cortex at this stage (Rice et al., 2012). It will be interesting to determine if these innate immune regulators play a role in regulating this pathway.

In this and previous studies, we have found that deletion of *ric8a* gene from radial glial progenitors using *nestin-cre* does not result in obvious cortical ectopia (**Fig. 2**) (Ma et al., 2017). However, when *ric8a* is in addition deleted from microglia, this results in severe cortical ectopia (**Fig. 2**), implicating a novel role of microglia in cortical ectopia development. Previous studies have reported that *ric8a* deletion by *nestin-cre* alone results in cortical ectopia (Kask et al., 2015; Kask et al., 2018). The cause for this discrepancy is at present unclear. The expression of *nestin-cre*, however, is known to be influenced by several factors including transgene insertion site and genetic background and the same *nestin-cre* has been reported to be active and induce gene inactivation in microglia (Karasinska et al., 2013; Takamori et al., 2009). These factors may play a role in this discrepancy. In our studies, we show that microglia specific *ric8a* deletion using *cx3cr1-cre* during development results in severe cortical ectopia upon and only upon immune stimulation (**Fig. 2**). We further show that microglia specific *app* deletion results in similar ectopia also only upon immune stimulation (**Fig. 3**). These results are important findings as they implicate, for the first time, a causative role played by microglial dysfunction in the formation of cortical ectopia in neurodevelopmental disorders.

### Neuronal activity, glial cytokine expression, and brain circuit plasticity

Activity-dependent competitive and homeostatic plasticity is a foundational rule that regulates the development, maturation, and function of neural circuits across brain regions. Studies have shown that glial cells, through regulating the spatiotemporal expression of immune cytokines, play a pivotal role in this process. In the developing thalamus, by activating interleukin-33 expression in an activity-dependent manner, astrocytes have been found to promote the segregation of eye-specific axonal projection and the maturation of the visual circuitry (He et al., 2022; Vainchtein et al., 2018). In the visual cortex, astrocytic expression of TNFα similarly mediates activity-dependent homeostatic upscaling of cortical synapses following peripheral monocular deprivation (Barnes et al., 2017; Heir et al., 2024; Kaneko et al., 2008). In this study, we have shown that Aβ monomers inhibit expression of cytokines by brain microglia via a novel APP/heterotrimeric G protein-mediated pathway. Aβ is primarily produced by neurons in the brain in a neural activity-dependent manner and form oligomers when large quantities are produced (Cirrito et al., 2005). Aβ oligomers, in contrary to monomers, are proinflammatory and increase glial cytokine expression (Halle et al., 2008; Huang, 2023; Lorton et al., 1996; Muehlhauser et al., 2001; Tan et al., 1999). These findings thus suggest that different levels of neural circuit activity in the brain may differentially regulate glial cytokine expression through inducing different levels of Aβ. High levels of neural activity may lead to high levels of Aβ and the formation of Aβ oligomers that activate glial cytokine production, while low levels of neural activity may produce low levels of Aβ, maintain Aβ as monomers, and inhibit glial cytokine production. Thus, Aβ in the brain may not only be a reporter of the levels of neural circuit activity but may also serve as an agent that directly mediate activity level-dependent plasticity. Following sensory deprivation, for example, Aβ levels may be lowered due to loss of sensory stimulation. This may lead to the relief of monomeric Aβ inhibition of cytokines such as TNFα and as a result trigger homeostatic upscaling of cortical synapses in the visual cortex (Barnes et al., 2017; Heir et al., 2024; Kaneko et al., 2008). In contrary, when neural activity levels are high, large quantities of Aβ may be produced, leading to formation of Aβ oligomers that may in turn induce expression of cytokines such as IL-33 that promote synaptic pruning. A large body of evidence strongly indicates that Aβ and related pathways indeed mediate homeostatic and competitive plasticity in the visual and other systems of the brain (Galanis et al., 2021; Huang, 2023, 2024; Kamenetz et al., 2003; Kim et al., 2013). Our discovery of the Aβ monomer-activated pathway has therefore provided novel insights into a universal mechanism that senses neural circuit activity pattern and translates it into homeostatic and competitive synaptic changes in the brain, a mechanism with fundamental roles in cognitive function.

In this study, we have also found that the matrix proteinase MMP9 is a key downstream effector of microglial activity in the developing cortex. We find that microglial hyperactivity results in increased levels of MMP9, leading to cortical basement membrane degradation and neuronal ectopia and inhibiting MMP9 directly or indirectly suppresses the phenotype. This suggests that the regulation of MMP9 may be a key mechanism by which glial cells regulate brain development and plasticity. Indeed, independent studies have shown that, in the visual cortex, MMP9 is also a pivotal mediator of TNFα-dependent homeostatic upscaling of central synapses following monocular deprivation (Akol et al., 2022; Kaneko et al., 2008; Kelly et al., 2015; Spolidoro et al., 2012). In the Xenopus tectum, MMP9 has similarly been found to be induced by neural activity and promote visual activity-induced dendritic growth (Gore et al., 2021). Importantly, in both wildtype and amblyopic animals, light reintroduction after dark exposure has been found to reactivate plasticity in the adult visual cortex via MMP9, uncovering a potential treatment for common visual conditions (Murase et al., 2017; Murase et al., 2019). These results therefore highlight a conserved glia/cytokine/MMP9-mediated mechanism that regulates brain development and plasticity from embryogenesis to adulthood. In ocular dominance plasticity, MMP9 is activated at perisynaptic regions (Murase et al., 2017; Murase et al., 2019). MMP9 mRNA translation has been also observed in dendrites (Dziembowska et al., 2012). In the *ric8a* mutant cortex, we find that MMP9 activity is increased. Further studies are required to precisely determine the cellular sources of MMP9 and how its activity is regulated.

### Aβ monomer anti-inflammatory activity and Alzheimer’s disease

Aβ is well known as a component of the amyloid plaques in the Alzheimer’s disease brain. It is a unique amphipathic peptide that can, dependent on concentration and other conditions, remain as monomers or form oligomers. Studies on Aβ have historically focused on the neurotoxic effects of Aβ oligomers and their proinflammatory effects on glia (Gulisano et al., 2018; Halle et al., 2008; He et al., 2019; Huang, 2023; Kim et al., 2013; Lauren et al., 2009; Lazarevic et al., 2017; Lorton et al., 1996; Muehlhauser et al., 2001; Parodi et al., 2010; Puzzo et al., 2008; Shankar et al., 2008; Tan et al., 1999; Walsh et al., 2002; Yang et al., 2015; Zott et al., 2019). In this study, we have found that, in contrary to Aβ oligomers, Aβ monomers instead possess a previously unknown anti-inflammatory activity that acts through a unique microglial pathway. We have further found that genetic disruption of this pathway in corticogenesis results microglial hyperactivity, leading to neuronal ectopia and large disruption of cortical structural organization. To our knowledge, ours is the first study to uncover this overlooked anti-inflammatory activity of Aβ monomers. It is in alignment with recent studies showing that Aβ monomers are also directly protective to neurons and positively regulate synapse development and function (Galanis et al., 2021; Giuffrida et al., 2009; Plant et al., 2003; Ramsden et al., 2002; Zhou et al., 2022). Assuming a set amount of Aβ peptides, the formation of Aβ oligomers and aggregates in the brain would, by chemical law, be predicted to result in the depletion of Aβ monomers (Dear et al., 2020; Michaels et al., 2020). Thus, in the Alzheimer’s disease brain, besides the obvious formation of Aβ aggregates, there may also be a less visible depletion of Aβ monomers taking place at the same time, which may, like Aβ oligomers, also contribute to the development of neuroinflammation and neuronal damage (Huang, 2023). In support of this interpretation, high soluble brain Aβ42, which likely also means high levels of Aβ monomers in the brain, have been found in clinical studies to preserve cognition in patients of both familial and sporadic Alzheimer’s disease, in spite of increasing amyloidosis detected in their brains (Espay et al., 2021; Sturchio et al., 2022; Sturchio et al., 2021). In our study, we have also found that the effects of microglial disinhibition are mediated by MMP9. Importantly, in neurodegenerative diseases, MMP9 has been similarly found to be a key determinant regulatings the selective degeneration of neuronal cell types (Kaplan et al., 2014; Tran et al., 2019). MMP9 levels are also upregulated in the plasma in both mild cognitive impairment and Alzheimer’s disease patients (Bruno et al., 2009; Lorenzl et al., 2008; Tsiknia et al., 2022). In addition, in several motor neuron disease models, reducing MMP9 has been found to protect neurons and delay the loss of motor function (Kaplan et al., 2014; Spiller et al., 2019). Thus, our study has not only uncovered a potentially overlooked role of Aβ monomer depletion in the development of Alzheimer’s disease but also identified downstream effectors. Elucidating the roles these factors play may reveal new insight into the pathogenesis of Alzheimer’s disease.

## METHODS

### Generation of *ric8a* conditional allele

Standard molecular biology techniques were employed for generating the conditional *ric-8a* allele. Briefly, genomic fragments, of 4.5 and 2.5 kilobases and flanking exons 2-4 of the *ric-8a* locus at the 5’ and 3’ side, respectively, were isolated by PCR using high fidelity polymerases. Targeting plasmid was constructed by flanking the genomic fragment containing exons 2-4 with two loxP sites together with a *neomycin* positive selection cassette, followed by 5’ and 3’ genomic fragments as homologous recombination arms and a *pgk-DTA* gene as a negative selection cassette. ES cell clones were screened by Southern blot analysis using external probes at 5’ and 3’ sides. For derivation of conditional allele, the *neomycin* cassette was removed by crossing to an *actin-flpe* transgenic line after blastocyst injection and germ line transmission. The primer set for genotyping *ric-8a* conditional allele, which produces a wildtype band of ∼110bp and a mutant band of ∼200bp, is: 5’-cctagttgtgaatcagaagcacttg-3’ and 5’-gccatacctgagttacctaggc-3’. Animals homozygous for the conditional *ric-8a* allele are viable and fertile, without obvious phenotypes.

### Mouse breeding and pharmacology

*emx1-cre, nestin*-*cre, foxg1-cre, cx3cr1-cre, floxed app* as well as the *BAT-lacZ* reporter mouse lines were purchased from the Jackson Lab. *nex-cre* and *wnt3a-cre* were as published (Goebbels et al., 2006; Yoshida et al., 2006). *cre* transgenes were introduced individually into the *ric8a or app* conditional mutant background for phenotypic analyses and *ric8a or app* homozygotes without *cre* as well as heterozygotes with *cre* (littermates) were both analyzed as controls. For BB94 and MMP9/13 inhibitor injection, pregnant females were treated daily from E12.5 to E14.5 at 30 μg (BB94) or 37.5 μg (MMP9/13 inhibitor) per g of body weight. For dorsomorphin and S3I-201 injection, pregnant females were treated on E12.5 at 7.5 and 25 μg per g of body weight, respectively. For sham treatment, pregnant females were treated on E12.5 with 100 μls of DMSO. BrdU was injected at 100 μg per g of body weight, and embryos were collected 4 hours later for cell proliferation analysis, or alternatively, pups were sacrificed at P5 for neuronal migration analysis and at P17 for other analysis. For LPS treatment, pregnant females were injected intraperitoneally with 400ng (*ric8a* genetic background) or 150ng (*app* genetic background) LPS per g of body weight on both E11.5 and E12.5. Animal use was in accordance with institutional guidelines.

### Immunohistochemistry

Vibratome sections from brains fixed in 4% paraformaldehyde were used. The following primary antibodies were used at respective dilutions/concentrations: mouse anti-BrdU supernatant (clone G3G4, Developmental Studies Hybridoma Bank (DSHB), University of Iowa, IA; 1:40), mouse anti-RC2 supernatant (DSHB; 1:10), mouse anti-Nestin supernatant (DSHB; 1:20), mouse anti-Vimentin supernatant (DSHB; 1:10), mouse anti-Pax6 supernatant (DSHB; 1:20), moue anti-Reelin (Millipore, 1:500), mouse anti-chondroitin sulfate (CS-56, Sigma, 1:100), rat anti-Ctip2 (Abcam, 1:500), rabbit anti-phospho Histone H3 (Ser10) (Millipore; 1:400), rabbit anti-Cux1 (CDP) (Santa Cruz; 1:100), rabbit anti-laminin (Sigma; 1:2000), rabbit anti-GFAP (Dako;1:1000), rabbit anti-ALDH1L1 (Abcam, 1:500), rabbit anti-MMP9 (Abcam, 1:1000), goat anti-MMP2 (R&D Systems; 5 μg/ml), rabbit anti-Calretinin (Chemicon, 1:2000), mouse anti-S100β (Thermo Scientific; 1:100), rabbit anti-S100β (Thermo Scientific; 1:200), and rabbit anti-phospho-Smad1/5 (Ser463/465) (41D10; Cell Signaling, 1:200). FITC and Cy3 conjugated secondary antibodies were purchased from Jackson ImmunoResearch Laboratories (West Grove, PA). Peroxidase conjugated secondary antibodies were purchased from Santa Cruz Biotech. Staining procedures were performed as described previously (Huang et al., 2006), except for anti-Ric-8a, MMP9, and phospho-Smad1/5 staining, in which a tyramide signal amplification (TSA) plus Cy3 kit (PerkinElmer, Waltham, MA) was used per manufacturer’s instruction. Sections were mounted with Fluoromount G medium (Southern Biotech, Birmingham, AB) and analyzed under a Nikon *eclipse* Ti microscope or an Olympus confocal microscope.

### Microglia culture and assay

Cerebral hemispheres were dissected from individual neonates, mechanically dissociated, split into 3-4 wells each and cultured in DMEM-F12 (Lonza) containing 10% fetal bovine serum (FBS) (Invitrogen). Microglial cells were harvested by light trypsinization that removes astroglial sheet on day 13-15. For experiments other than assaying IL-1β secretion, microglia were treated with LPS at 20ng/ml for 3 hours or at 5ng/ml overnight and, if applicable, DMSO or Aβ40 (ApexBio and Genscript) was applied at the same time as LPS. For assaying IL-1β secretion, microglia were primed with LPS at 200ng/ml for 5-6 hours before treatment with 3mM ATP for 15 minutes. In these experiments, DMSO or Aβ40 was applied at the same time as ATP if applicable. Supernatants were collected and used for cytokine ELISA assays per manufacturer’s instructions (Biolegend). Total RNAs were prepared from collected cells using Trizol (Invitrogen) and cDNAs were synthesized using a High-capacity cDNA reverse transcription kit (Applied Biosystems). Quantitative PCR was performed using a GoTaq qPCR master mix per manufacturer’s instructions (Promega). All gene expression levels were normalized against that of GAPDH.

### Quantitative and statistical analysis

The sample size was estimated to be 3-9 animals each genotype (every 4^th^ of 50 μm coronal sections, 7-10 sections each animal) for ectopia analysis, 3-5 animals each genotype (3-4 sections each animal) for immunohistochemical analysis, and 4-6 animals each genotype for gel zymography and Western blot analysis, as has been demonstrated by previous publications to be adequate for similar animal studies. Matching sections were used between controls and mutants. NIS-Elements BR 3.0 software (Nikon) was used for quantifying the numbers and sizes of neuronal ectopia, the numbers of laminin positive debris, as well as the numbers of astrocytes. ImageJ software (NIH) was used for quantifying the intensity of immunostainings. In analysis of radial glial cell division, the cleavage plane angle was calculated by determining the angle between the equatorial plate and the ventricular surface. Statistics was performed using Student’s *t* test when comparing two conditions, or one-way ANOVA followed by Tukey’s post hoc test when comparing three or more conditions. All data are represented as means ± s.e.m.

## ACKNOWLEDGMENTS

1. Z. H. thanks Dr. L. F. Reichardt for supporting the initial generation of *ric8a* mutant ES cells, Dr. E. A. Grove (Chicago) for providing the *wnt3a-cre* strain, the late Dr. B. A. Barres (Stanford) for critiques and input, and Drs. W. L. Murphy and E. Bresnick (UW-Madison) for access to a plate reader and a qPCR machine. We also thank the late Dr. D. Oertel (UW-Madison) for critical reading and editing and Dr. L Puglielli (UW-Madison) for critical reading of a previous version of the manuscript. This work was supported by funds from the Departments of Neurology and Neuroscience, UW-Madison, and a Basil O’Connor award from the March of Dimes foundation to Z.H.

## AUTHOR INFORMATION

Z. H. designed experiments, generated *ric8a* conditional ES cells, performed microglial and related experiments, and wrote the manuscript. H. J. K., D. S., and Z. H. performed other experiments and analyzed data.

---

**Supplemental Fig. 1.**
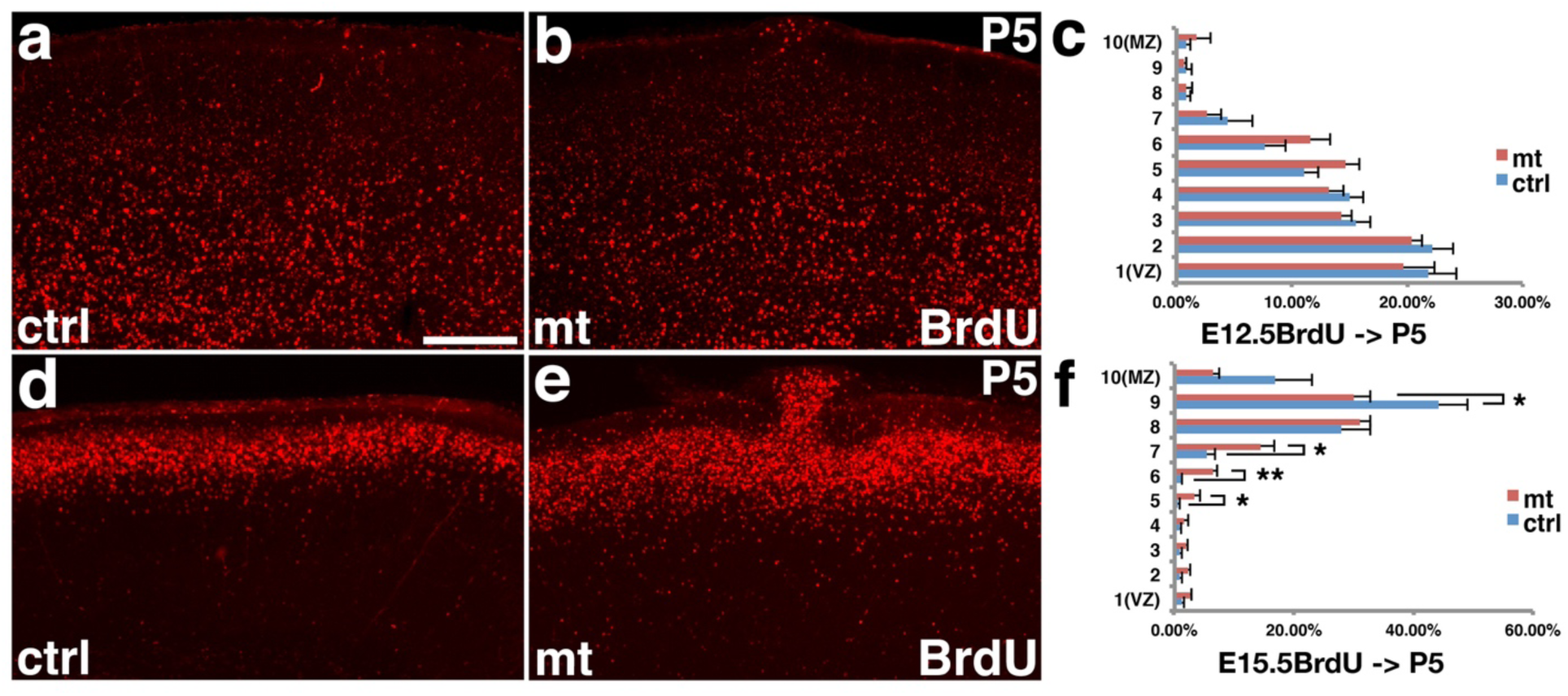
Birth-dating of early and late-born neurons in *ric8a-emx1-cre* mutant cortices. (**a-c**) BrdU (in red) staining in control (**a**) and mutant (**b**) cortices at P5 after administration at E12.5. Quantification is shown in (**c**). No statistically significant differences were observed between control and mutant neurons in regions without ectopia. (**d-f**) BrdU staining in control (**d**) and mutant (**e**) cortices at P5 after administration at E15.5. Quantification is shown in (**f**). Neuronal migration appears slightly delayed in mutants as compared to controls. *, *P* < 0.05; **, *P* < 0.01; n = 5.

**Supplemental Fig. 2.**
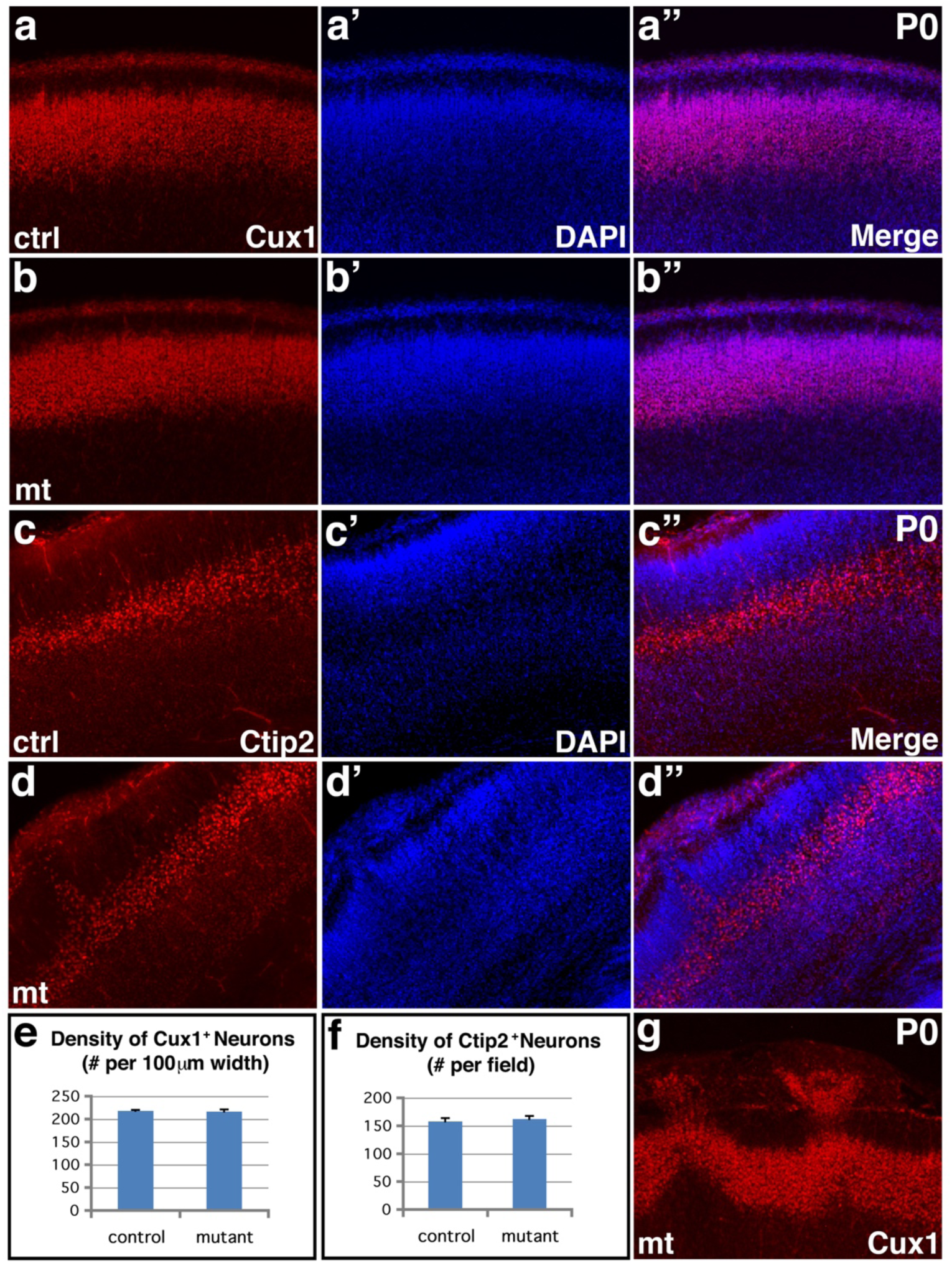
Lamina-specific neuronal markers are normal outside ectopia in *ric8a*-*emx1-cre* mutant cortices. **(a**-**b**”) Cux1 (in red) and nuclear (DAPI, in blue) staining of control (**a-a”**) and mutant (**b-b”**) cortices at P0 in a region without ectopia. No obvious changes in the expression pattern of Cux1, an upper layer neuronal marker, were observed in the mutant cortex, except in areas with ectopia (see panel (**g**)). (**c**-**d**”) Ctip2 (in red) and nuclear (DAPI, in blue) staining of control (**c**-**c**”) and mutant (**d**-**d**”) cortices at P0. No obvious changes in the expression pattern of Ctip2, a deep layer neuronal marker, were observed in the mutant cortex, except in areas with ectopia. (**e-f**) Quantification of cortical neurons positive for Cux1 (**e**) and Ctip2 (**f**) in matching cortical regions at P0. No significant differences were observed in the density of Cux1 (control, 218.1 ± 1.7 per 100 μm cortical width; mutant, 216.4 ± 4.3 per 100 μm cortical width; *P* = 0.36, n = 12) or Ctip2 (control, 157.8 ± 5.0 per field; mutant, 161.9 ± 5.9 per field; *P* = 0.31, n = 12) positive neurons between controls and mutants. (**g**) Cux1 (in red) staining of mutant cortices at P0 in a region with ectopia. Scale bar in (**a**), 200 μm for all panels.

**Supplemental Fig. 3.**
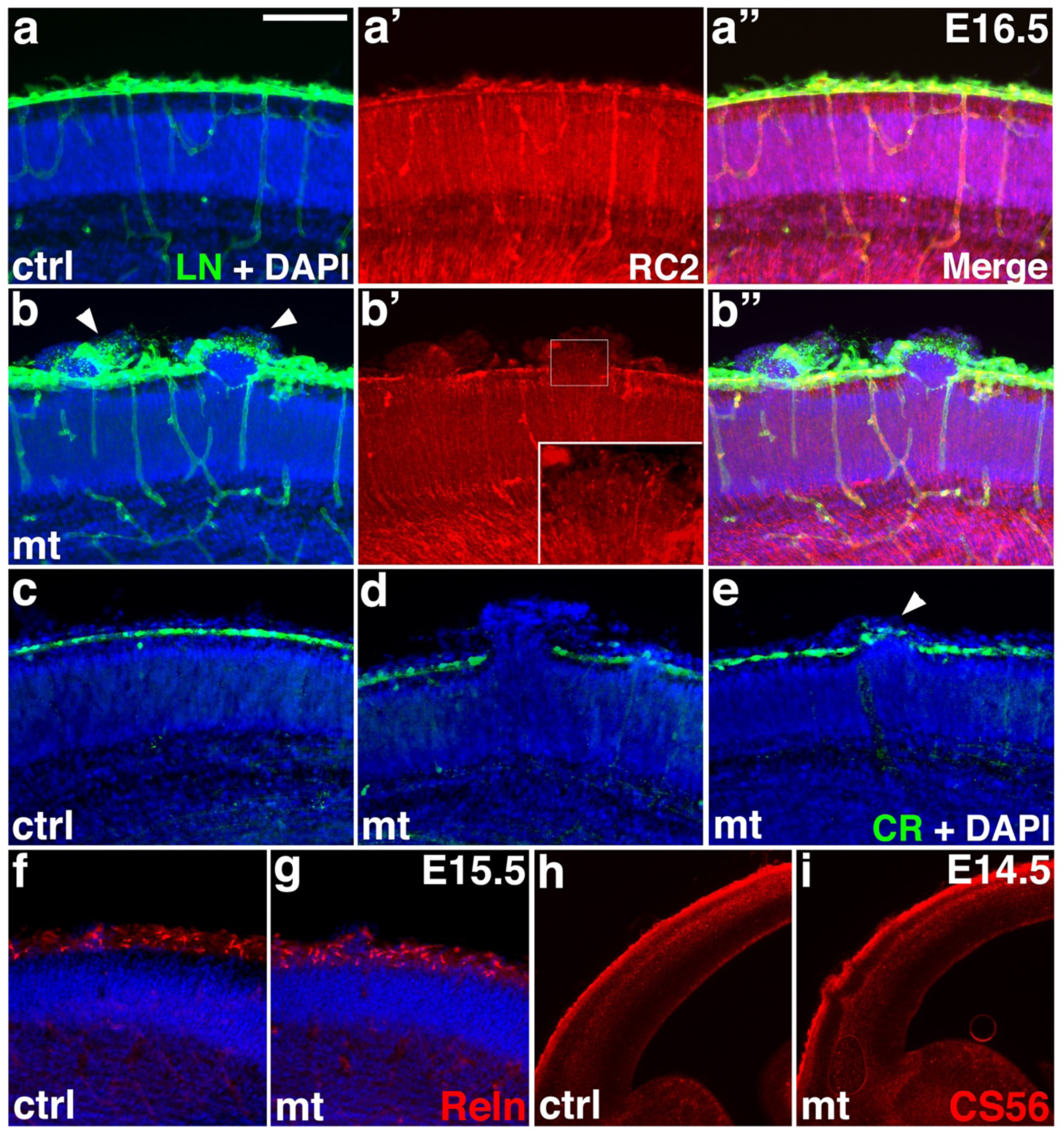
Neuronal ectopia in *ric8a-emx1-cre* mutants result from pial basement membrane breach during embryogenesis. **(a**-**a”**) Laminin (LN, in green), radial glial marker RC2 (in red), and nuclear (DAPI, in blue) staining of control cortices at E16.5. A continuous basement membrane is observed at the pia, where radial glial endfeet are anchored. (**b**-**b”**) Laminin, RC2, and nuclear staining of *ric8a/emx1-cre* mutant cortices at E16.5. Neuronal ectopias are consistently observed at sites of basement membrane breakage (arrowheads in **b**). Radial glial fibers at these sites extend beyond the pia (inset in **b’**). (**c**-**e**) Calretinin (CR, in green) and nuclear (DAPI, in blue) staining of control (**c**) and mutant (**d-e**) cortices at E16.5. A continuous row of Calretinin positive Cajal-Retzius cells is observed in the marginal zone of control cortices (**c**). By contrast, in mutants, Cajal-Retzius cells are absent at large ectopias (**d**). However, they appear passively displaced by over-migrating neurons at small ectopias (arrowhead in **e**). **(f-g**) Reelin (Reln, in red) and nuclear (DAPI, in blue) staining of control (**f**) and *ric8a*/*emx1-cre* mutant (**g**) cortices at E15.5. Strong Reelin expression is observed in Cajal-Retzius cells in the marginal zone of both control and mutant cortices. **(h-i)** Chondroitin sulfate proteoglycan (CS56, in red) staining of control (**h**) and mutant (**i**) cortices at E14.5. Normal preplate splitting is observed in mutants. Scale bar in (**a**), 200μm for (**a**-**g)** and 500μm for (**h**-**i**).

**Supplemental Fig. 4.**
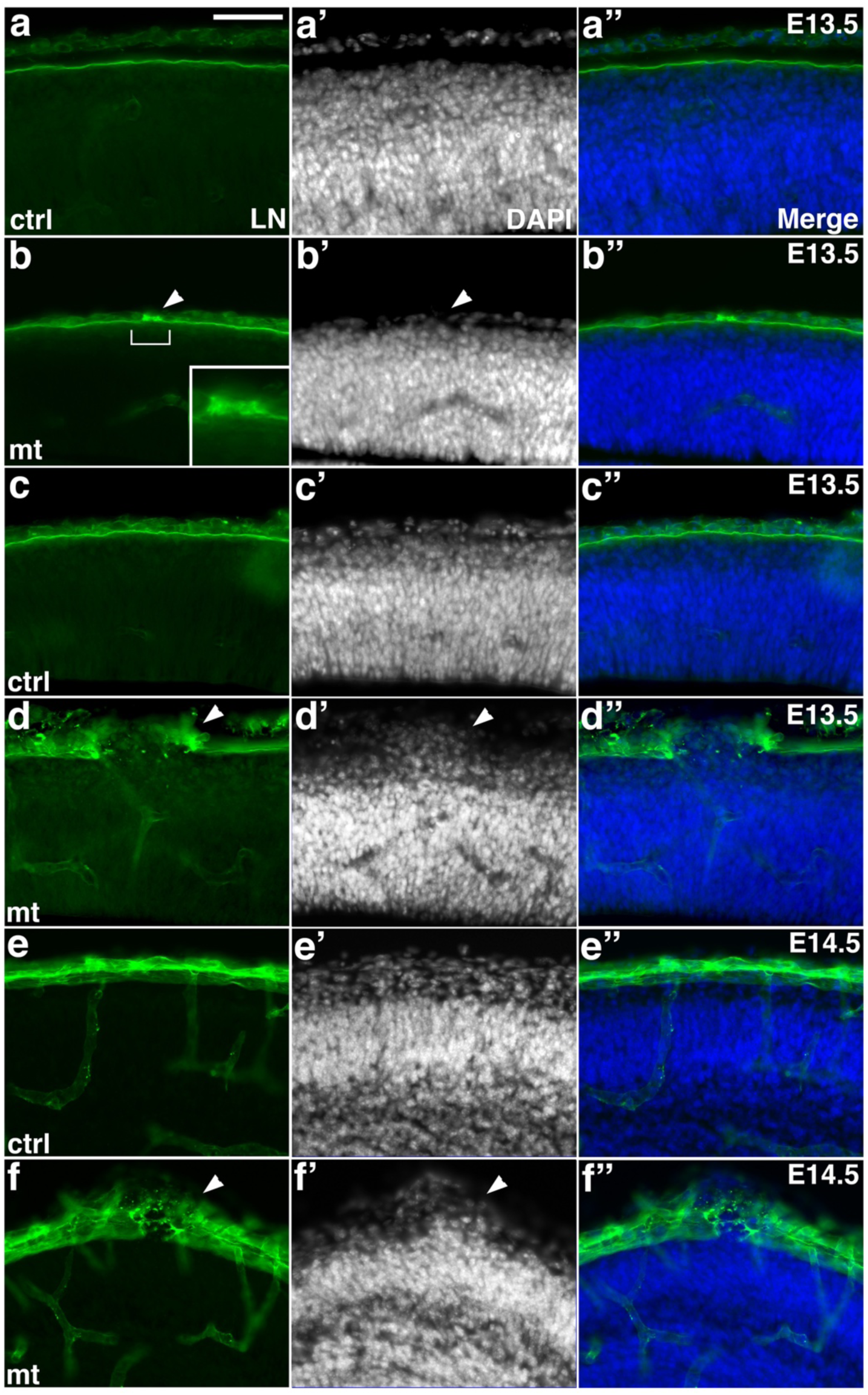
Basement membrane breaches precede neuronal ectopia in *ric8a*-*emx1-cre* mutant cortices. **(a**-**a”**) Laminin (LN, in green) and nuclear (DAPI, in blue) staining of control cortices at E13.5. A continuous basement membrane is observed at the pia, beneath which cells are well organized in the cortical wall. (**b**-**b”**) Laminin and nuclear staining of *ric8a*-*emx1-cre* mutant cortices at E13.5. In a subset of mutants, a small disruption of basement membrane is observed (bracket and inset in **b**), but not yet associated with ectopia (arrowhead in **b’**). **(c**-**c”**) Laminin (LN, in green) and nuclear (DAPI, in blue) staining of control cortices at E13.5. (**d**-**d”**) Laminin and nuclear staining of *ric8a*-*emx1-cre* mutant cortices at E13.5. Although at E13.5 we observe basement membrane defects in the absence of neuronal ectopia (see **b-b”**), when there are neuronal ectopia, they are always associated with basement membrane breakage. (**e**-**e”**) Laminin and nuclear staining of control cortices at E14.5. (**f**-**f”**) Laminin and nuclear staining of *ric-8a-emx1-cre* mutant cortices at E14.5. Neuronal ectopia at E14.5 are also always associated with basement membrane breakage. Scale bar in (**a**), 100 μm for all panels.

**Supplemental Fig. 5.**
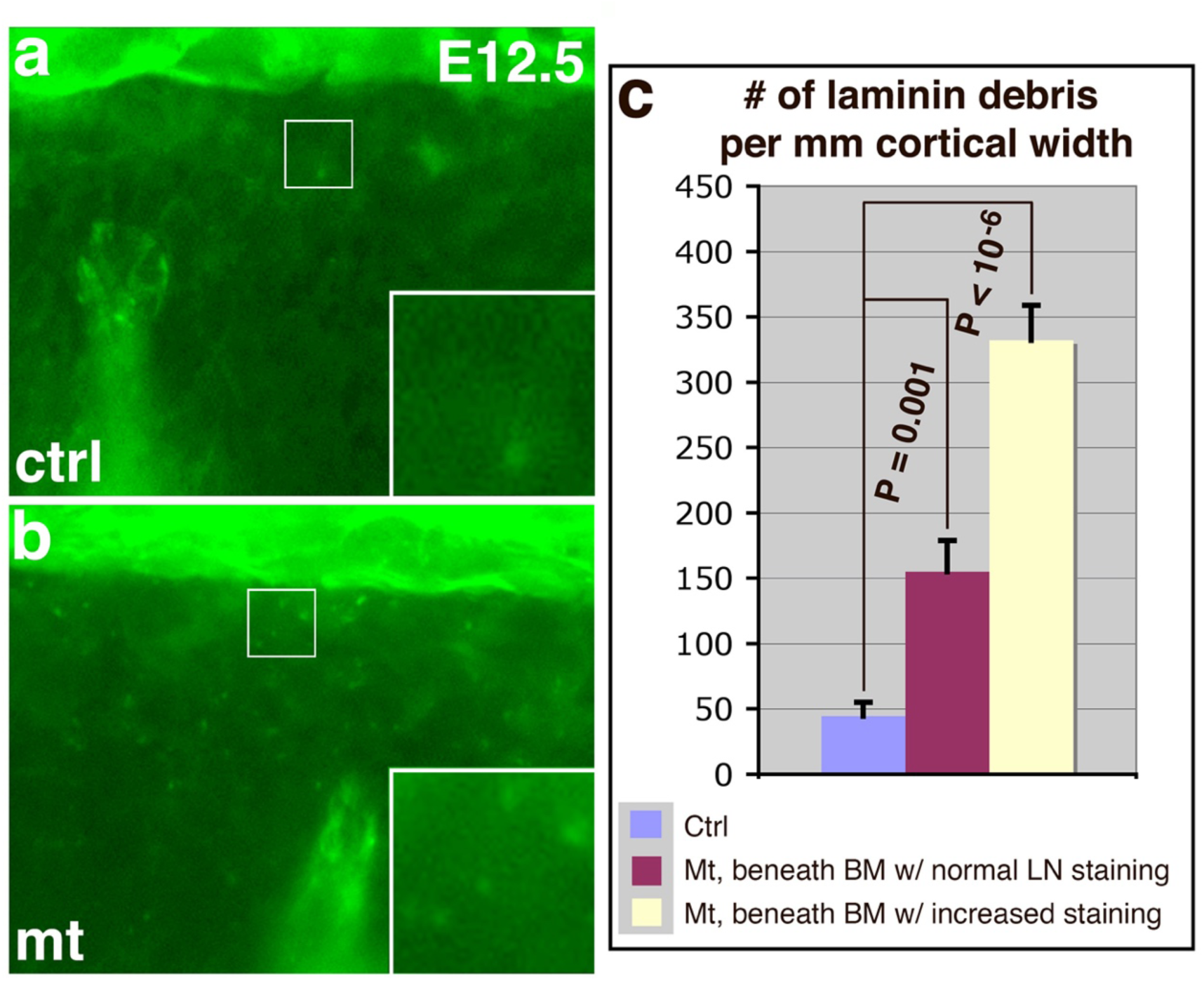
Signs of basement membrane degradation before breach formation at E12.5. (**a-b**) Laminin (in green) staining of control (**a**) and *ric8a-emx1-cre* mutant (**b**) cortices at E12.5. Increased numbers of laminin positive debris were observed in mutants (compare insets), even though breaches had yet to form. (**c**) Quantitative analysis shows significant increases.

**Supplemental Fig. 6.**
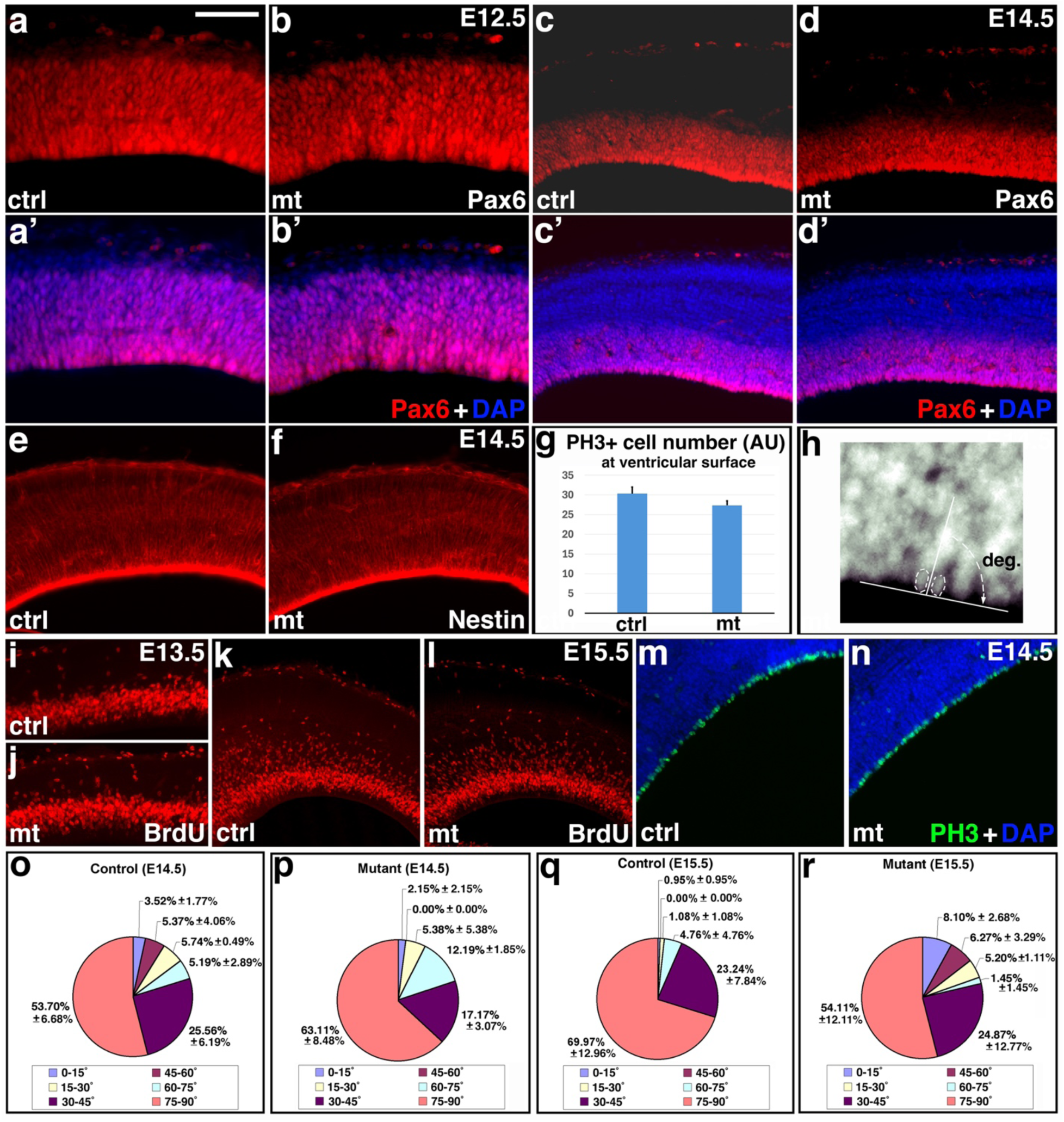
Cortical radial glial identity and proliferation are unaffected in *ric8a*-*emx1-cre* mutants. **(a**-**b’**) Pax6 (in red) and nuclear (DAPI, in blue) staining of control (**a**&**a’**) and mutant (**b**&**b’**) cortices at E12.5. (**c**-**d’**) Pax6 (in red) and nuclear (DAPI, in blue) staining of control (**c**&**c’**) and mutant (**d**&**d’**) cortices at E14.5. No ectopic Pax6 positive cells were observed at either E12.5 or E14.5. **(e-f**) Nestin (in red) and nuclear (DAPI, in blue) staining of control (**e**) and mutant (**f**) cortices at E14.5. (g) Quantification showed no significant differences in the number of phospho-histone 3 (PH3) positive cells at the ventricular surface between control and mutants at E14.5 (AU, arbitrary units; *P* = 0.15, n= 9 each). See also images in (**m-n**). (h) Cleavage plane of neural progenitors is defined by the angle between the equatorial plate and the ventricular surface. See quantification results in (**o-r**). **(i-j**) BrdU staining (in red) in control (**i**) and mutant (**j**) cortices at E13.5. (**k-l**) BrdU staining (in red) in control (**k**) and mutant (**l**) cortices at E15.5. (**m-n**) Phospho-histone 3 (PH3, in green) and nuclear (DAPI, in blue) staining of control (**m**) and mutant (**n**) cortices at E14.5. **(o-p**) Cleavage plane distribution of radial glial mitosis in control (**o**) and mutant (**p**) cortices at E14.5. No significant differences were observed (*P* > 0.4, n = 3 animals each genotype; 73 cells for controls and 76 cells for mutants). (**q**-**r**) Cleavage plane distribution of radial glial mitosis in control (**q**) and mutant (**r**) cortices at E15.5. No significant differences were observed (*P* > 0.1, n = 3 animals each genotype; 70 cells for controls and 59 cells for mutants). Scale bar in (**a**), 100 μm for (**a**-**b’**) and 200 μm for (**c-n**).

**Supplemental Fig. 7.**
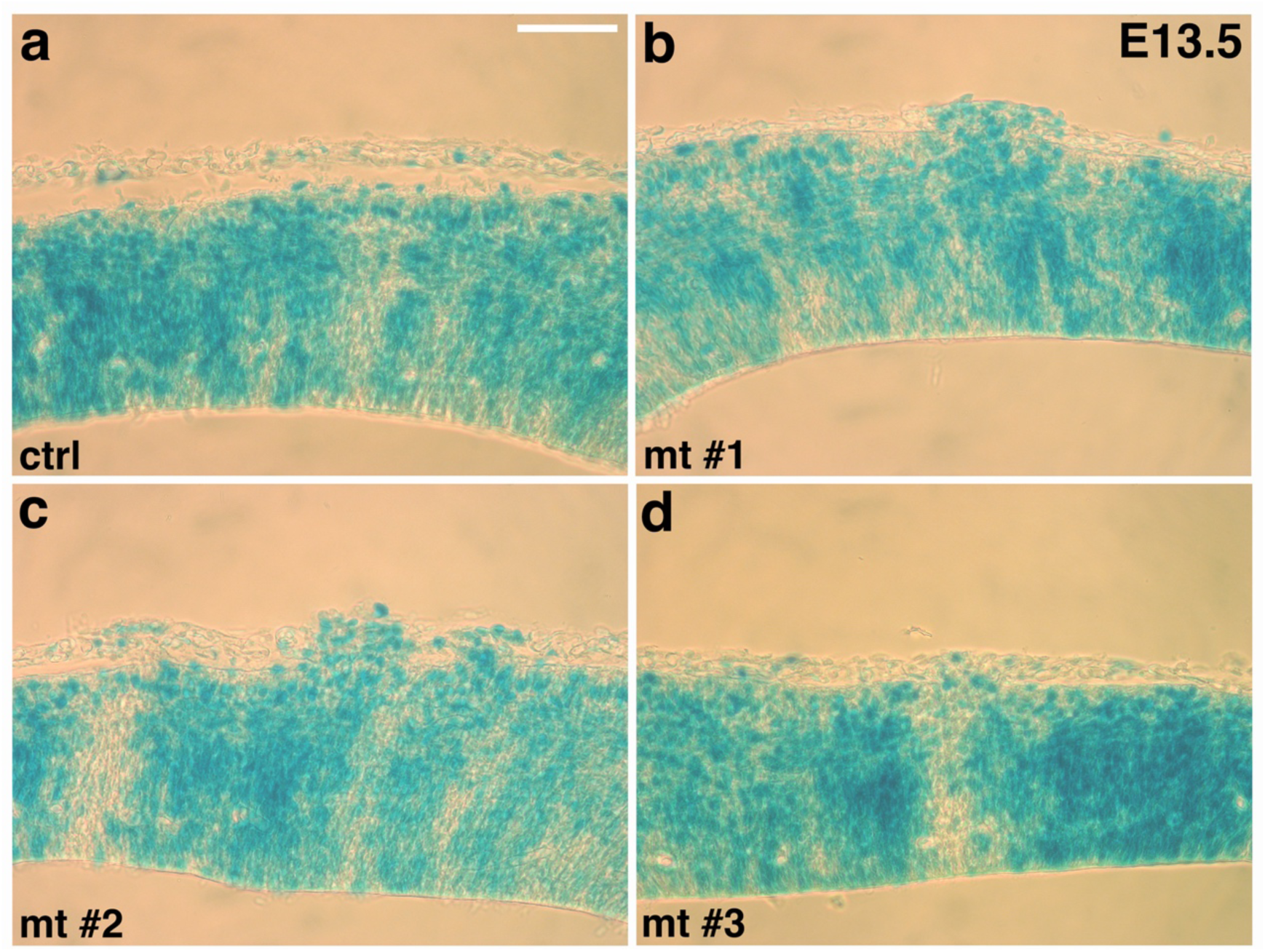
Wnt pathway activity is normal in *ric8a*-*emx1-cre* mutant cortices. X-gal staining of BAT-lacZ expression in *ric8a*-*emx1-cre* control (**a**) and mutant (**b**-**d**) cortices at E13.5. No obvious differences are observed between controls and three different mutants at this stage.

**Supplemental Fig. 8.**
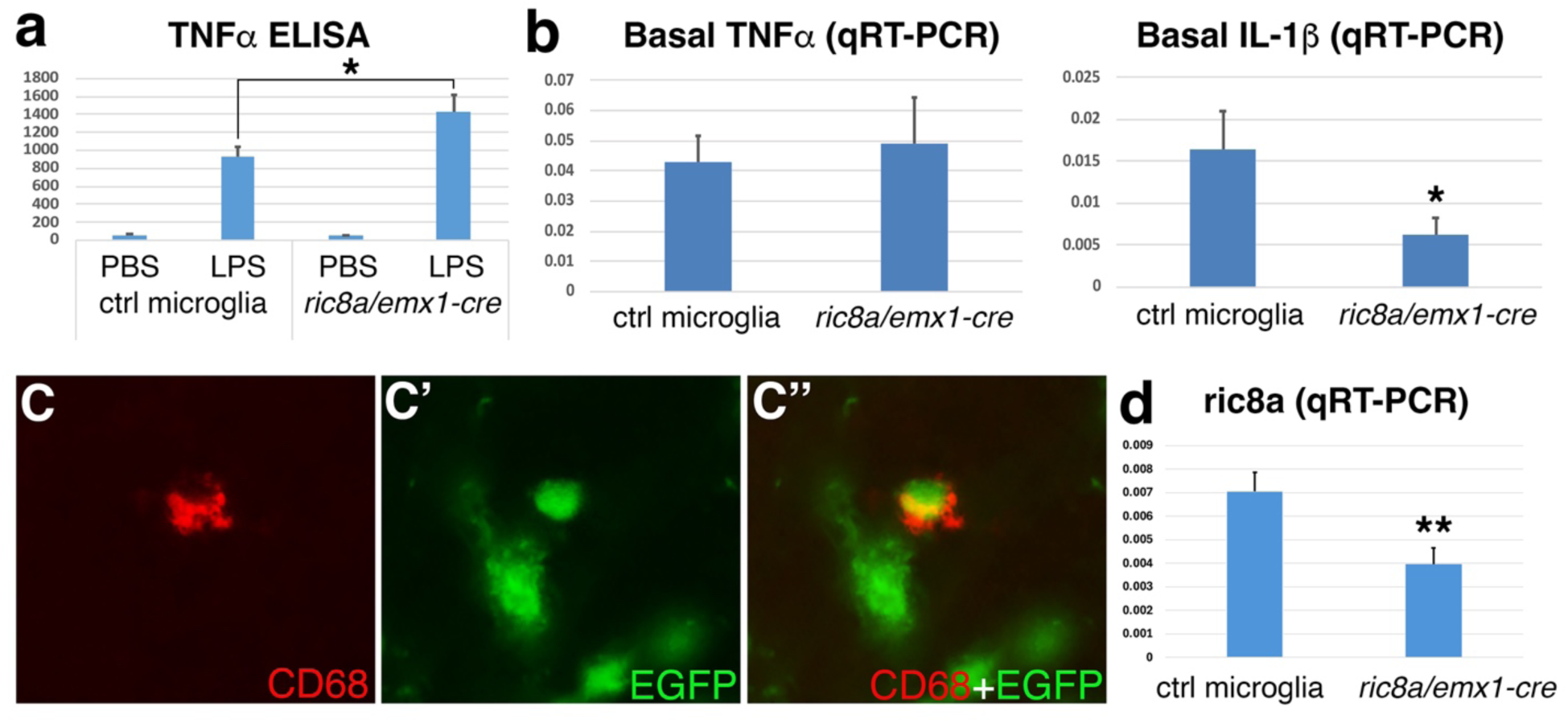
*emx1-cre* is active in microglia. (**a-b**) TNFα secretion (pg/ml) (**a**) and basal TNFα and IL-1β mRNA expression (**b**) in control and *ric8a-emx1-cre* mutant microglia. *, *P* < 0.05; n = 5-8 each group. (**c-c”**) The Rosa26 EGFP (green) is induced in CD68 positive (red) microglial cells in *ric8a-emx1-cre* mutant cortices. (**d**) Quantitative RT-PCR analysis of microglia cultured from *ric8a-emx1-cre* mutant cortices showed severe loss of *ric8a* mRNA in microglial cells. **, *P* < 0.01; n = 5-8 each group.

**Supplemental Fig. 9.**
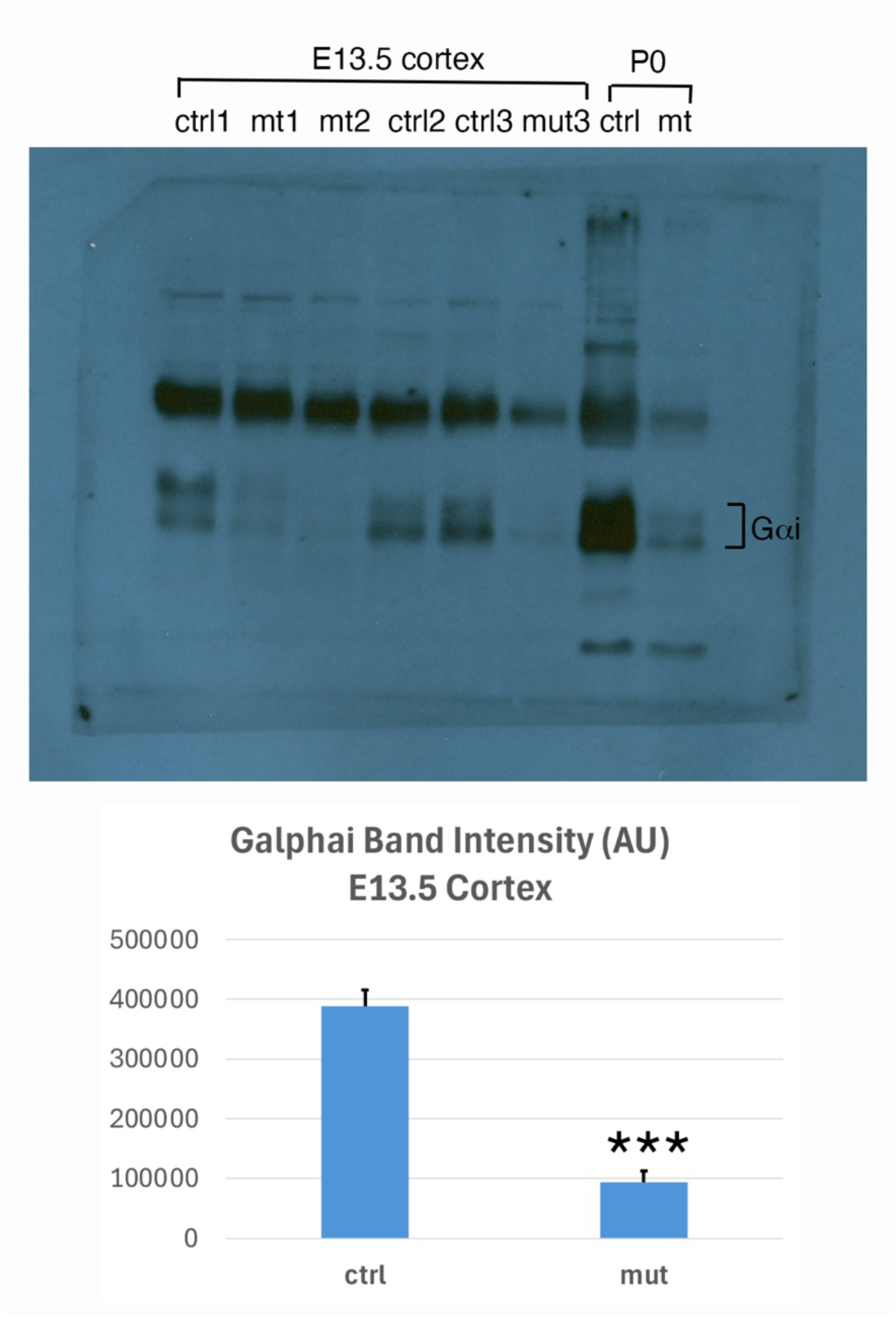
Gαi protein is severely depleted from *ric8a-emx1-cre* mutant cortices. Western blot analysis of Gai proteins in E13.5 *ric8a-emx1-cre* mutant cortices showed that Gai protein levels were severely reduced (AU, arbitrary units). *P* < 0.001, n = 3 each.

**Supplemental Fig. 10.**
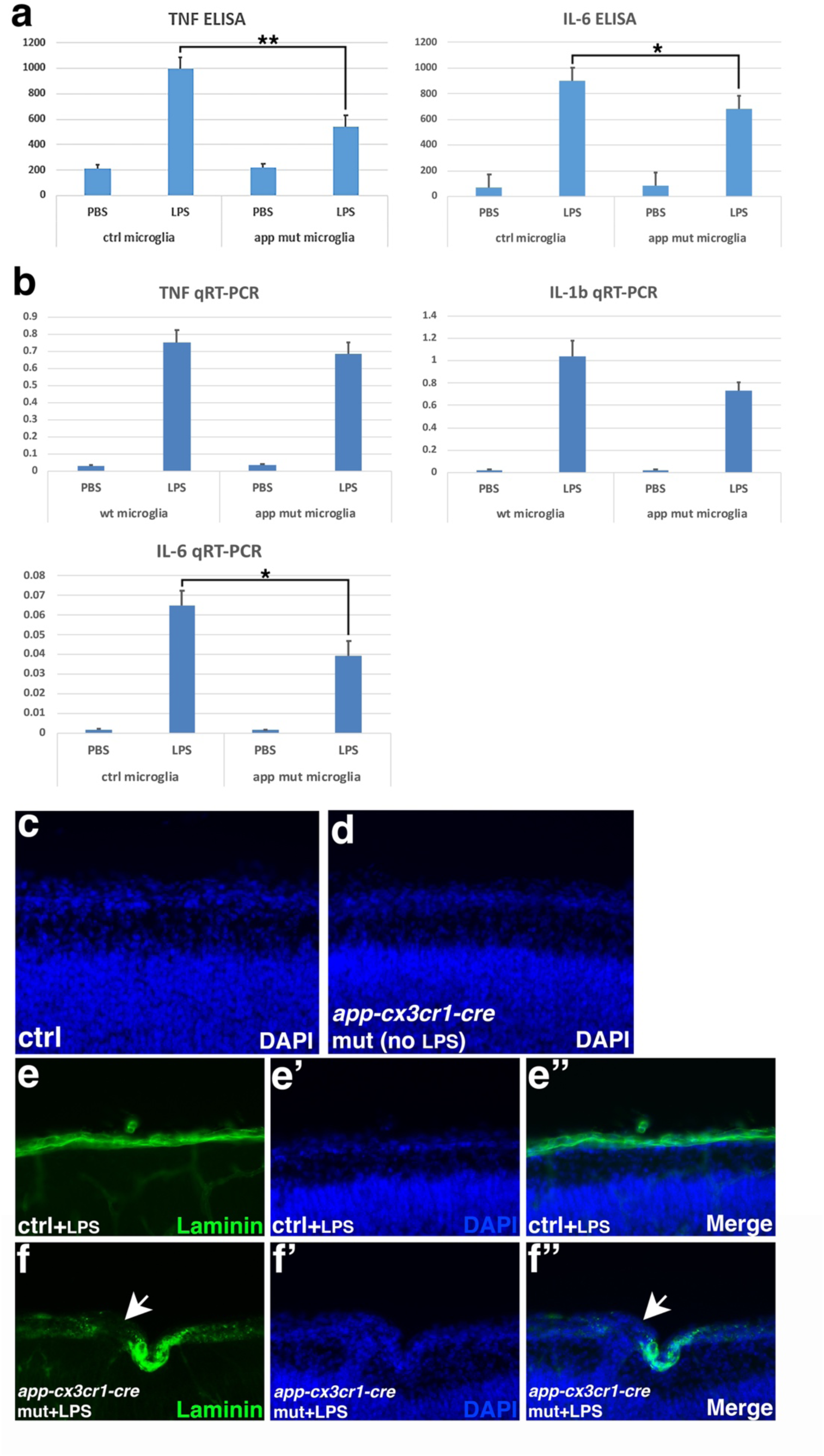
Cytokine secretion and transcriptional induction in *app-cx3cr1-cre* mutant microglia. **(a)** TNFα and IL-6 secretion (pg/ml) in control and *app-cx3cr1-cre* mutant microglia following overnight LPS stimulation. *, *P* < 0.05; **, *P* < 0.01; n = 9-13 each group. **(b)** TNFα, IL-1β and IL-6 mRNA expression in control and *app-cx3cr1-cre* mutant microglia following overnight 3-hr LPS stimulation. *, *P* < 0.05; n = 6-7 each group. (**c-d**) *app-cx3cr1-cre* mutant cortices showed no ectopia at P0 without LPS treatment at embryonic stages (DAPI, blue) (**e-f”**) *app-cx3cr1-cre* mutant cortices treated with LPS at embryonic stages showed perturbed basement membrane and gaps at sites of ectopia (white arrows) at P0 (Laminin, green; DAPI, blue).

**Supplemental Fig. 11.**
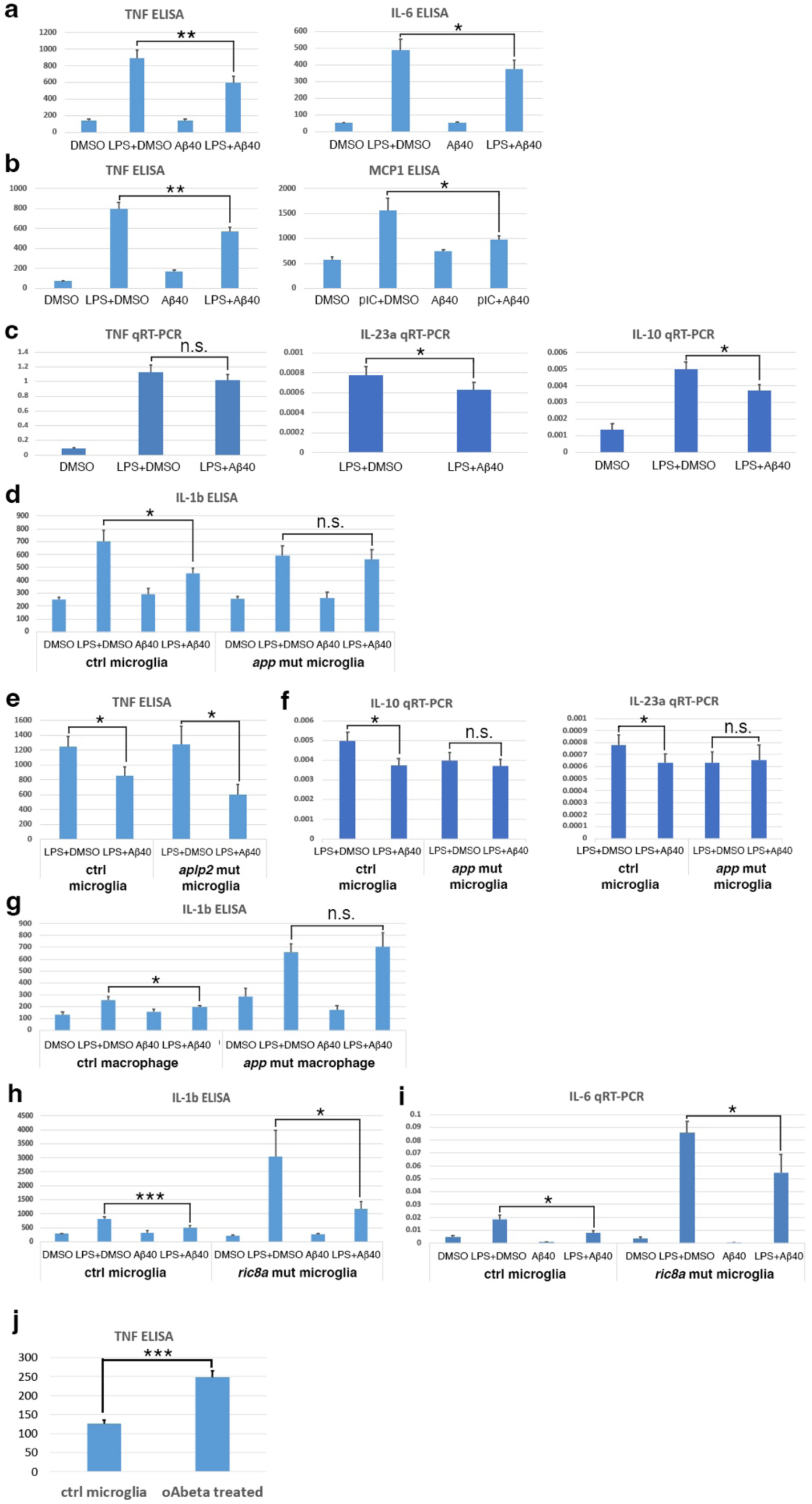
Effects of monomeric Aβ on cytokine secretion and transcription in control and mutant microglial lineage cells. **(a)** TNFα and IL-6 secretion (pg/ml) in wildtype microglia following LPS stimulation in the absence or presence of Aβ40 (50nM). *, *P* < 0.05; **, *P* < 0.01; n = 25 each group for TNFα and 11 each group for IL-6. **(b)** TNFα and MCP1 secretion (pg/ml) in wildtype microglia following LPS stimulation in the absence or presence of Aβ40 (500nM) from Genscript. Effects on IL-1β secretion in Fig. 4b was also performed with Genscript Aβ40. All other experiments in Fig. 7 were performed with ApexBio Aβ40. *, *P* < 0.05; **, *P* < 0.01; n = 5-7 each group. **(c)** TNFα, IL-23, and IL-10 mRNA expression in wildtype microglia following LPS stimulation in the absence or presence of Aβ40 (400nM). *, *P* < 0.05; n = 6 each group **(d)** IL-1β secretion (pg/ml) in control and *app/cx3cr1-cre* mutant microglia following LPS stimulation in the absence or presence of Aβ40. *, *P* < 0.05; n = 8-12 each group. **(e)** TNFα (pg/ml) in control or *aplp2/cx3cr1-cre* mutant microglia following LPS stimulation in the absence or presence of Aβ40 (400nM). *, *P* < 0.05; n = 9-13 each group. **(f)** IL-10 and IL-23 mRNA expression in control and *app/cx3cr1-cre* mutant microglia following LPS stimulation in the absence or presence of Aβ40 (400nM). *, *P* < 0.05; n = 6 each group. **(g)** IL-1β secretion (pg/ml) in fresh unelicited control and *app/cx3cr1-cre* mutant peritoneal macrophages following LPS stimulation in the absence or presence of Aβ40 (400nM). *, *P* < 0.05; n = 12 each group. **(h)** IL-1β secretion (pg/ml) in f control and *ric8a/cx3cr1-cre* mutant microglia following LPS stimulation in the absence or presence of Aβ40 (500nM). *, *P* < 0.05; ***, *P* < 0.001; n = 7-8 each group. **(i)** IL-6 mRNA expression in control and *ric8a/cx3cr1-cre* mutant microglia following LPS stimulation in the absence or presence of Aβ40 (200nM). *, *P* < 0.05; n = 6 each group. **(j)** TNFα (pg/ml) in wildtype microglia in the absence or presence of Aβ40 oligomers aggregated (at 10μM monomer equivalent). ***, *P* < 0.001; n = 10 each group.

**Supplemental Fig. 12.**
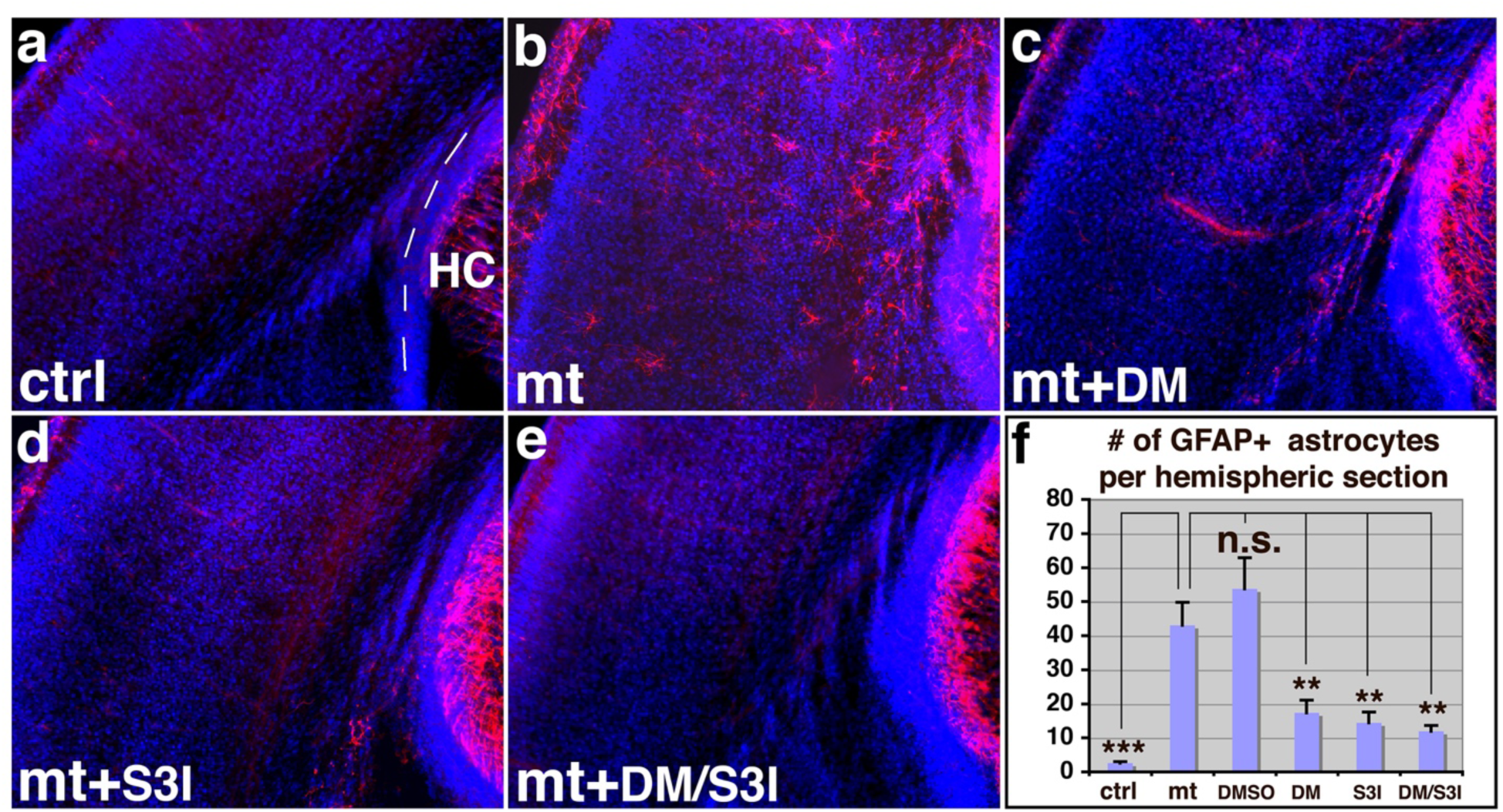
Suppression of astrogliosis in *ric8a*-*emx1-cre* mutant cortices by anti-inflammatory drugs, dorsomorphin (DM) and S3I-201 (S3I). **(a-e**) GFAP (in red) and nuclear (DAPI, in blue) staining of neonatal control (**a**) and mutant cortices without treatment (**b**) or mutant cortices after dorsomorphin (DM, **c**), S3I-201 (S3I, **d**), or dual (DM+S3I, **e**) treatment at E12.5. Note, GFAP is normally expressed in the neonatal hippocampus (HC) (dashed line in (**a**)). **(f**) Quantitative analysis of GFAP-positive astrocyte numbers in the neonatal mutant cortex after treatment at E12.5.

**Supplemental Fig. 13.**
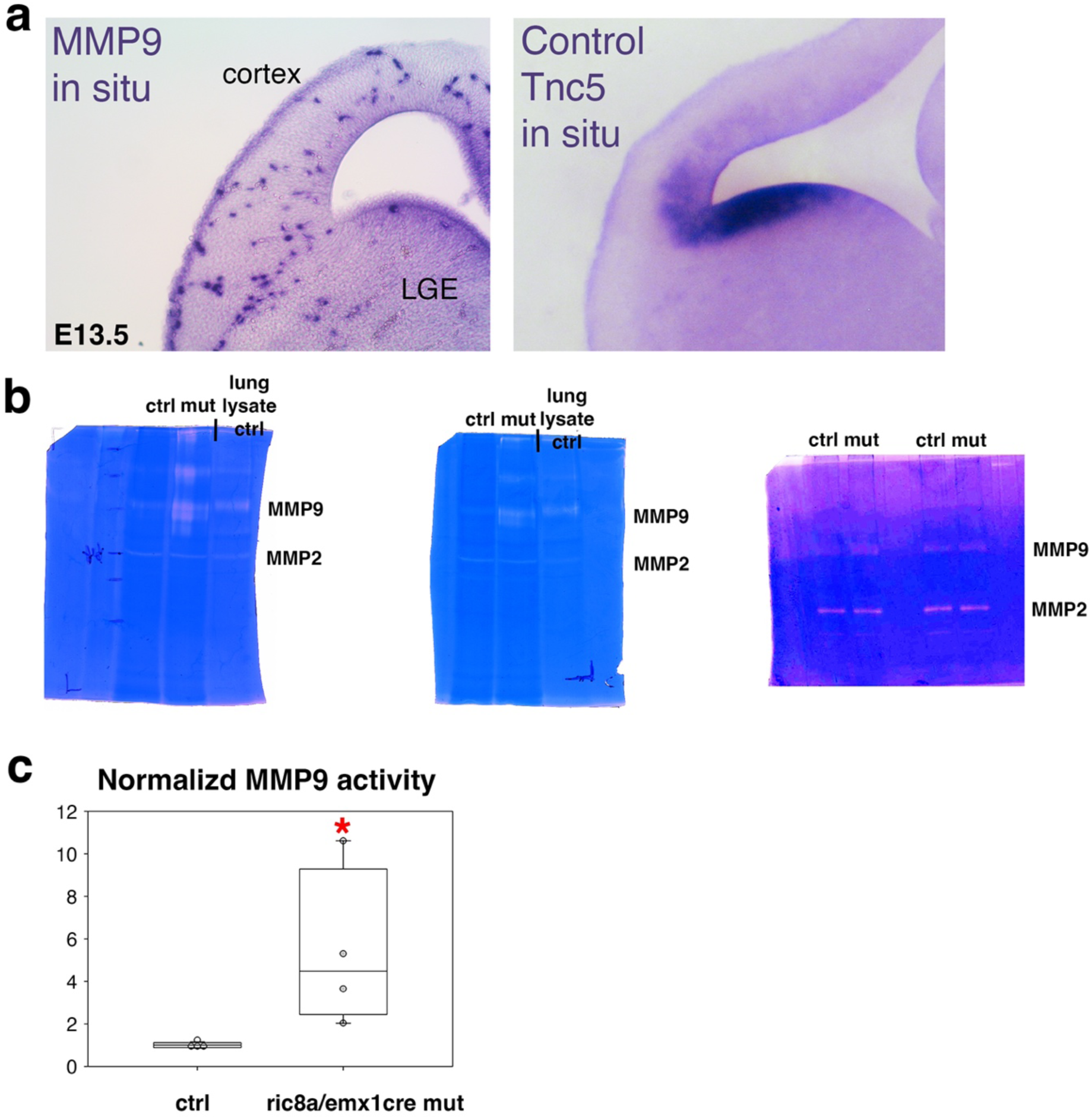
MMP9 in situ and activity in E13.5 *ric8a*-*emx1-cre* mutant cortices. (a) Sections of E13.5 brain wholemount in situ of MMP9 showed a sparse MMP9 expressing cell population resembling microglia (LGE, lateral ganglionic eminence). Tnc5 in situ was performed as control for validating probe specificity. (b) Whole gel gelatin zymography images of *ric8a*-*emx1-cre* control and mutant cortices. Embryonic lung lysates were used as control for validation of MMP2/9 activity. (c) Quantification showed MMP9 activity levels were significantly increased in E13.5 *, *P* < 0.05; n = 4 each.

**Supplemental Fig. 14.**
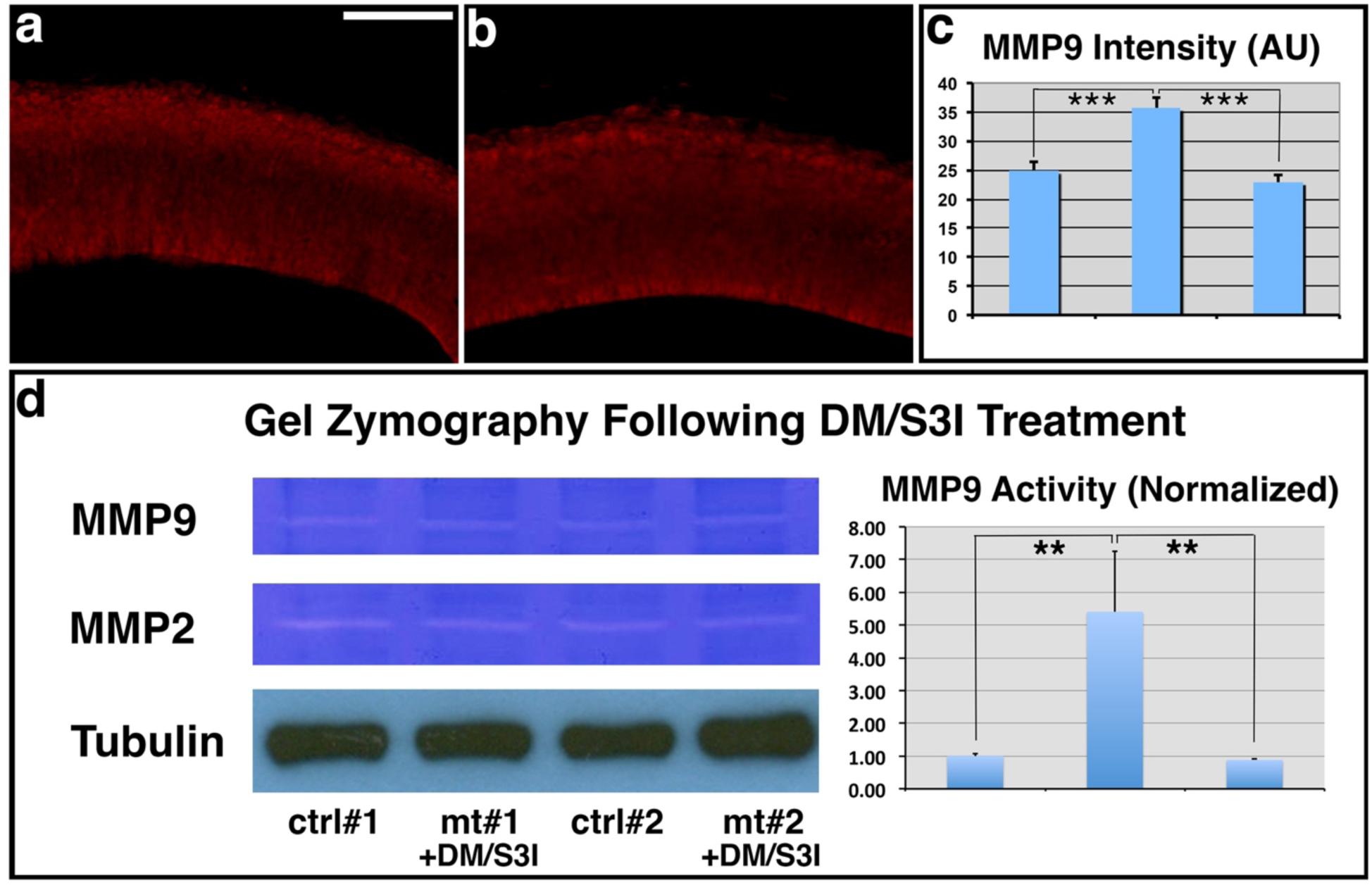
Suppression of MMP9 expression in *ric8a*-*emx1-cre* mutant cortices by anti-inflammatory drugs, dorsomorphin (DM) and S3I-201 (S3I). **(a)** MMP9 (in red) staining in control cortices at E13.5. **(b)** MMP9 (in red) staining in mutant cortices at E13.5 after DM and S3I dual treatment at E12.5. **(c)** Quantitative analysis of MMP9 expression. No significant differences are observed in mutants after inhibitor treatment in comparison to controls (***, *P* < 0.001; n = 6 each group, ANOVA). **(d)** Gel zymography of E13.5 control and mutant cortical lysates following DM/S3I treatment at E12.5. Similar levels of MMP9 are observed between controls and mutants. Quantification also showed no significant differences in normalized MMP9 levels (**, *P* < 0.01; n = 4-6 each group, ANOVA). See also supplemental Fig.13b-c.

## Notes

### Competing Interest Statement

The authors have declared no competing interest.

### Summary of Updates

Supp Figs 8, 10, 11, 14 revised; New Supp Figs. 9, 13

## REFERENCES

1. Afshar, K., Willard, F.S., Colombo, K., Johnston, C.A., McCudden, C.R., Siderovski, D.P., and Gonczy, P. (2004). RIC-8 is required for GPR-1/2-dependent Galpha function during asymmetric division of C. elegans embryos. Cell 119, 219–230.

2. Akol, I., Kalogeraki, E., Pielecka-Fortuna, J., Fricke, M., and Lowel, S. (2022). MMP2 and MMP9 Activity Is Crucial for Adult Visual Cortex Plasticity in Healthy and Stroke-Affected Mice. The Journal of neuroscience : the official journal of the Society for Neuroscience 42, 16–32.

3. Amaya, D.A., Wegner, M., Stolt, C.C., Chehrehasa, F., Ekberg, J.A., and St John, J.A. (2015). Radial glia phagocytose axonal debris from degenerating overextending axons in the developing olfactory bulb. J Comp Neurol 523, 183–196.

4. Barnes, S.J., Franzoni, E., Jacobsen, R.I., Erdelyi, F., Szabo, G., Clopath, C., Keller, G.B., and Keck, T. (2017). Deprivation-Induced Homeostatic Spine Scaling In Vivo Is Localized to Dendritic Branches that Have Undergone Recent Spine Loss. Neuron 96, 871–882 e875.

5. Barres, B.A. (2008). The mystery and magic of glia: a perspective on their roles in health and disease. Neuron 60, 430–440.

6. Beggs, H.E., Schahin-Reed, D., Zang, K., Goebbels, S., Nave, K.A., Gorski, J., Jones, K.R., Sretavan, D., and Reichardt, L.F. (2003). FAK deficiency in cells contributing to the basal lamina results in cortical abnormalities resembling congenital muscular dystrophies. Neuron 40, 501–514.

7. Belvindrah, R., Nalbant, P., Ding, S., Wu, C., Bokoch, G.M., and Muller, U. (2006). Integrin-linked kinase regulates Bergmann glial differentiation during cerebellar development. Molecular and cellular neurosciences 33, 109–125.

8. Billings, E.A., Lee, C.S., Owen, K.A., D’Souza, R.S., Ravichandran, K.S., and Casanova, J.E. (2016). The adhesion GPCR BAI1 mediates macrophage ROS production and microbicidal activity against Gram-negative bacteria. Science signaling 9, ra14.

9. Blaschke, A.J., Staley, K., and Chun, J. (1996). Widespread programmed cell death in proliferative and postmitotic regions of the fetal cerebral cortex. Development 122, 1165–1174.

10. Blaschke, A.J., Weiner, J.A., and Chun, J. (1998). Programmed cell death is a universal feature of embryonic and postnatal neuroproliferative regions throughout the central nervous system. J Comp Neurol 396, 39–50.

11. Bruno, M.A., Mufson, E.J., Wuu, J., and Cuello, A.C. (2009). Increased matrix metalloproteinase 9 activity in mild cognitive impairment. J Neuropathol Exp Neurol 68, 1309–1318.

12. Chamberlain, L.M., Holt-Casper, D., Gonzalez-Juarrero, M., and Grainger, D.W. (2015). Extended culture of macrophages from different sources and maturation results in a common M2 phenotype. J Biomed Mater Res A 103, 2864–2874.

13. Cirrito, J.R., Yamada, K.A., Finn, M.B., Sloviter, R.S., Bales, K.R., May, P.C., Schoepp, D.D., Paul, S.M., Mennerick, S., and Holtzman, D.M. (2005). Synaptic activity regulates interstitial fluid amyloid-beta levels in vivo. Neuron 48, 913–922.

14. Colonna, M., and Butovsky, O. (2017). Microglia Function in the Central Nervous System During Health and Neurodegeneration. Annu Rev Immunol 35, 441–468.

15. Couwenbergs, C., Spilker, A.C., and Gotta, M. (2004). Control of embryonic spindle positioning and Galpha activity by C. elegans RIC-8. Current biology : CB 14, 1871–1876.

16. Cunningham, C.L., Martinez-Cerdeno, V., and Noctor, S.C. (2013). Microglia regulate the number of neural precursor cells in the developing cerebral cortex. The Journal of neuroscience : the official journal of the Society for Neuroscience 33, 4216–4233.

17. David, N.B., Martin, C.A., Segalen, M., Rosenfeld, F., Schweisguth, F., and Bellaiche, Y. (2005). Drosophila Ric-8 regulates Galphai cortical localization to promote Galphai-dependent planar orientation of the mitotic spindle during asymmetric cell division. Nature cell biology 7, 1083–1090.

18. Dear, A.J., Michaels, T.C.T., Meisl, G., Klenerman, D., Wu, S., Perrett, S., Linse, S., Dobson, C.M., and Knowles, T.P.J. (2020). Kinetic diversity of amyloid oligomers. Proceedings of the National Academy of Sciences of the United States of America 117, 12087–12094.

19. Dziembowska, M., Milek, J., Janusz, A., Rejmak, E., Romanowska, E., Gorkiewicz, T., Tiron, A., Bramham, C.R., and Kaczmarek, L. (2012). Activity-dependent local translation of matrix metalloproteinase-9. The Journal of neuroscience : the official journal of the Society for Neuroscience 32, 14538–14547.

20. Eichler, A., Kleidonas, D., Turi, Z., Fliegauf, M., Kirsch, M., Pfeifer, D., Masuda, T., Prinz, M., Lenz, M., and Vlachos, A. (2023). Microglial Cytokines Mediate Plasticity Induced by 10 Hz Repetitive Magnetic Stimulation. The Journal of neuroscience : the official journal of the Society for Neuroscience 43, 3042–3060.

21. Espay, A.J., Sturchio, A., Schneider, L.S., and Ezzat, K. (2021). Soluble Amyloid-beta Consumption in Alzheimer’s Disease. J Alzheimers Dis 82, 1403–1415.

22. Flak, M.B., Koenis, D.S., Sobrino, A., Smith, J., Pistorius, K., Palmas, F., and Dalli, J. (2020). GPR101 mediates the pro-resolving actions of RvD5n-3 DPA in arthritis and infections. J Clin Invest 130, 359–373.

23. Fogel, H., Frere, S., Segev, O., Bharill, S., Shapira, I., Gazit, N., O’Malley, T., Slomowitz, E., Berdichevsky, Y., Walsh, D.M., et al. (2014). APP homodimers transduce an amyloid-beta-mediated increase in release probability at excitatory synapses. Cell reports 7, 1560–1576.

24. Gabay, M., Pinter, M.E., Wright, F.A., Chan, P., Murphy, A.J., Valenzuela, D.M., Yancopoulos, G.D., and Tall, G.G. (2011). Ric-8 proteins are molecular chaperones that direct nascent G protein alpha subunit membrane association. Science signaling 4, ra79.

25. Galanis, C., Fellenz, M., Becker, D., Bold, C., Lichtenthaler, S.F., Muller, U.C., Deller, T., and Vlachos, A. (2021). Amyloid-Beta Mediates Homeostatic Synaptic Plasticity. The Journal of neuroscience : the official journal of the Society for Neuroscience 41, 5157–5172.

26. Garcia-Osta, A., and Alberini, C.M. (2009). Amyloid beta mediates memory formation. Learn Mem 16, 267–272.

27. Ginhoux, F., Greter, M., Leboeuf, M., Nandi, S., See, P., Gokhan, S., Mehler, M.F., Conway, S.J., Ng, L.G., Stanley, E.R., et al. (2010). Fate mapping analysis reveals that adult microglia derive from primitive macrophages. Science 330, 841–845.

28. Ginisty, A., Gely-Pernot, A., Abaamrane, L., Morel, F., Arnault, P., Coronas, V., and Benzakour, O. (2015). Evidence for a subventricular zone neural stem cell phagocytic activity stimulated by the vitamin K-dependent factor protein S. Stem cells 33, 515–525.

29. Giuffrida, M.L., Caraci, F., Pignataro, B., Cataldo, S., De Bona, P., Bruno, V., Molinaro, G., Pappalardo, G., Messina, A., Palmigiano, A., et al. (2009). Beta-amyloid monomers are neuroprotective. The Journal of neuroscience : the official journal of the Society for Neuroscience 29, 10582–10587.

30. Goebbels, S., Bormuth, I., Bode, U., Hermanson, O., Schwab, M.H., and Nave, K.A. (2006). Genetic targeting of principal neurons in neocortex and hippocampus of NEX-Cre mice. Genesis 44, 611–621.

31. Gore, S.V., James, E.J., Huang, L.C., Park, J.J., Berghella, A., Thompson, A.C., Cline, H.T., and Aizenman, C.D. (2021). Role of matrix metalloproteinase-9 in neurodevelopmental deficits and experience-dependent plasticity in Xenopus laevis. eLife 10.

32. Gorski, J.A., Talley, T., Qiu, M., Puelles, L., Rubenstein, J.L., and Jones, K.R. (2002). Cortical excitatory neurons and glia, but not GABAergic neurons, are produced in the Emx1-expressing lineage. The Journal of neuroscience : the official journal of the Society for Neuroscience 22, 6309–6314.

33. Grant, J.L., Ghosn, E.E., Axtell, R.C., Herges, K., Kuipers, H.F., Woodling, N.S., Andreasson, K., Herzenberg, L.A., Herzenberg, L.A., and Steinman, L. (2012). Reversal of paralysis and reduced inflammation from peripheral administration of beta-amyloid in TH1 and TH17 versions of experimental autoimmune encephalomyelitis. Sci Transl Med 4, 145ra105.

34. Graus-Porta, D., Blaess, S., Senften, M., Littlewood-Evans, A., Damsky, C., Huang, Z., Orban, P., Klein, R., Schittny, J.C., and Muller, U. (2001). Beta1-class integrins regulate the development of laminae and folia in the cerebral and cerebellar cortex. Neuron 31, 367–379.

35. Guenette, S., Chang, Y., Hiesberger, T., Richardson, J.A., Eckman, C.B., Eckman, E.A., Hammer, R.E., and Herz, J. (2006). Essential roles for the FE65 amyloid precursor protein-interacting proteins in brain development. The EMBO journal 25, 420–431.

36. Gulisano, W., Melone, M., Li Puma, D.D., Tropea, M.R., Palmeri, A., Arancio, O., Grassi, C., Conti, F., and Puzzo, D. (2018). The effect of amyloid-beta peptide on synaptic plasticity and memory is influenced by different isoforms, concentrations, and aggregation status. Neurobiol Aging 71, 51–60.

37. Gulisano, W., Melone, M., Ripoli, C., Tropea, M.R., Li Puma, D.D., Giunta, S., Cocco, S., Marcotulli, D., Origlia, N., Palmeri, A., et al. (2019). Neuromodulatory Action of Picomolar Extracellular Abeta42 Oligomers on Presynaptic and Postsynaptic Mechanisms Underlying Synaptic Function and Memory. The Journal of neuroscience : the official journal of the Society for Neuroscience 39, 5986–6000.

38. Halle, A., Hornung, V., Petzold, G.C., Stewart, C.R., Monks, B.G., Reinheckel, T., Fitzgerald, K.A., Latz, E., Moore, K.J., and Golenbock, D.T. (2008). The NALP3 inflammasome is involved in the innate immune response to amyloid-beta. Nat Immunol 9, 857–865.

39. Hampoelz, B., Hoeller, O., Bowman, S.K., Dunican, D., and Knoblich, J.A. (2005). Drosophila Ric-8 is essential for plasma-membrane localization of heterotrimeric G proteins. Nat Cell Biol 7, 1099–1105.

40. Hattori, Y., Kato, D., Murayama, F., Koike, S., Asai, H., Yamasaki, A., Naito, Y., Kawaguchi, A., Konishi, H., Prinz, M., et al. (2023). CD206(+) macrophages transventricularly infiltrate the early embryonic cerebral wall to differentiate into microglia. Cell reports 42, 112092.

41. He, D., Xu, H., Zhang, H., Tang, R., Lan, Y., Xing, R., Li, S., Christian, E., Hou, Y., Lorello, P., et al. (2022). Disruption of the IL-33-ST2-AKT signaling axis impairs neurodevelopment by inhibiting microglial metabolic adaptation and phagocytic function. Immunity 55, 159–173 e159.

42. He, Y., Wei, M., Wu, Y., Qin, H., Li, W., Ma, X., Cheng, J., Ren, J., Shen, Y., Chen, Z., et al. (2019). Amyloid beta oligomers suppress excitatory transmitter release via presynaptic depletion of phosphatidylinositol-4,5-bisphosphate. Nat Commun 10, 1193.

43. Hebert, J.M., and McConnell, S.K. (2000). Targeting of cre to the Foxg1 (BF-1) locus mediates loxP recombination in the telencephalon and other developing head structures. Developmental biology 222, 296–306.

44. Heir, R., Abbasi, Z., Komal, P., Altimimi, H.F., Franquin, M., Moschou, D., Chambon, J., and Stellwagen, D. (2024). Astrocytes Are the Source of TNF Mediating Homeostatic Synaptic Plasticity. The Journal of neuroscience : the official journal of the Society for Neuroscience 44.

45. Herms, J., Anliker, B., Heber, S., Ring, S., Fuhrmann, M., Kretzschmar, H., Sisodia, S., and Muller, U. (2004). Cortical dysplasia resembling human type 2 lissencephaly in mice lacking all three APP family members. The EMBO journal 23, 4106–4115.

46. Huang, Z. (2023). A Function of Amyloid-beta in Mediating Activity-Dependent Axon/Synapse Competition May Unify Its Roles in Brain Physiology and Pathology. J Alzheimers Dis 92, 29–57.

47. Huang, Z. (2024). Evidence that Alzheimer’s Disease Is a Disease of Competitive Synaptic Plasticity Gone Awry. J Alzheimers Dis 99, 447–470.

48. Huang, Z., Shimazu, K., Woo, N.H., Zang, K., Muller, U., Lu, B., and Reichardt, L.F. (2006). Distinct roles of the beta 1-class integrins at the developing and the mature hippocampal excitatory synapse. J Neurosci 26, 11208–11219.

49. Iyer, H., Shen, K., Meireles, A.M., and Talbot, W.S. (2022). A lysosomal regulatory circuit essential for the development and function of microglia. Sci Adv 8, eabp8321.

50. Kamenetz, F., Tomita, T., Hsieh, H., Seabrook, G., Borchelt, D., Iwatsubo, T., Sisodia, S., and Malinow, R. (2003). APP processing and synaptic function. Neuron 37, 925–937.

51. Kaneko, M., Stellwagen, D., Malenka, R.C., and Stryker, M.P. (2008). Tumor necrosis factor-alpha mediates one component of competitive, experience-dependent plasticity in developing visual cortex. Neuron 58, 673–680.

52. Kaplan, A., Spiller, K.J., Towne, C., Kanning, K.C., Choe, G.T., Geber, A., Akay, T., Aebischer, P., and Henderson, C.E. (2014). Neuronal matrix metalloproteinase-9 is a determinant of selective neurodegeneration. Neuron 81, 333–348.

53. Karasinska, J.M., de Haan, W., Franciosi, S., Ruddle, P., Fan, J., Kruit, J.K., Stukas, S., Lutjohann, D., Gutmann, D.H., Wellington, C.L., et al. (2013). ABCA1 influences neuroinflammation and neuronal death. Neurobiol Dis 54, 445–455.

54. Kask, K., Ruisu, K., Tikker, L., Karis, K., Saare, M., Meier, R., Karis, A., Tonissoo, T., and Pooga, M. (2015). Deletion of RIC8A in neural precursor cells leads to altered neurogenesis and neonatal lethality of mouse. Dev Neurobiol 75, 984–1002.

55. Kask, K., Tikker, L., Ruisu, K., Lulla, S., Oja, E.M., Meier, R., Raid, R., Velling, T., Tonissoo, T., and Pooga, M. (2018). Targeted deletion of RIC8A in mouse neural precursor cells interferes with the development of the brain, eyes, and muscles. Dev Neurobiol 78, 374–390.

56. Kelly, E.A., Russo, A.S., Jackson, C.D., Lamantia, C.E., and Majewska, A.K. (2015). Proteolytic regulation of synaptic plasticity in the mouse primary visual cortex: analysis of matrix metalloproteinase 9 deficient mice. Front Cell Neurosci 9, 369.

57. Kim, T., Vidal, G.S., Djurisic, M., William, C.M., Birnbaum, M.E., Garcia, K.C., Hyman, B.T., and Shatz, C.J. (2013). Human LilrB2 is a beta-amyloid receptor and its murine homolog PirB regulates synaptic plasticity in an Alzheimer’s model. Science 341, 1399–1404.

58. Lauren, J., Gimbel, D.A., Nygaard, H.B., Gilbert, J.W., and Strittmatter, S.M. (2009). Cellular prion protein mediates impairment of synaptic plasticity by amyloid-beta oligomers. Nature 457, 1128–1132.

59. Lazarevic, V., Fienko, S., Andres-Alonso, M., Anni, D., Ivanova, D., Montenegro-Venegas, C., Gundelfinger, E.D., Cousin, M.A., and Fejtova, A. (2017). Physiological Concentrations of Amyloid Beta Regulate Recycling of Synaptic Vesicles via Alpha7 Acetylcholine Receptor and CDK5/Calcineurin Signaling. Front Mol Neurosci 10, 221.

60. Leathers, T.A., and Rogers, C.D. (2022). Time to go: neural crest cell epithelial-to-mesenchymal transition. Development 149.

61. Lee, Y., Park, B.H., and Bae, E.J. (2016). Compound C inhibits macrophage chemotaxis through an AMPK-independent mechanism. Biochemical and biophysical research communications 469, 515–520.

62. LeVine, H., 3rd (2004). Alzheimer’s beta-peptide oligomer formation at physiologic concentrations. Anal Biochem 335, 81–90.

63. Lewitus, G.M., Konefal, S.C., Greenhalgh, A.D., Pribiag, H., Augereau, K., and Stellwagen, D. (2016). Microglial TNF-alpha Suppresses Cocaine-Induced Plasticity and Behavioral Sensitization. Neuron 90, 483–491.

64. Liu, W., Yan, M., Liu, Y., Wang, R., Li, C., Deng, C., Singh, A., Coleman, W.G., Jr., and Rodgers, G.P. (2010). Olfactomedin 4 down-regulates innate immunity against Helicobacter pylori infection. Proceedings of the National Academy of Sciences of the United States of America 107, 11056–11061.

65. Lorenzl, S., Buerger, K., Hampel, H., and Beal, M.F. (2008). Profiles of matrix metalloproteinases and their inhibitors in plasma of patients with dementia. Int Psychogeriatr 20, 67–76.

66. Lorton, D., Kocsis, J.M., King, L., Madden, K., and Brunden, K.R. (1996). beta-Amyloid induces increased release of interleukin-1 beta from lipopolysaccharide-activated human monocytes. Journal of neuroimmunology 67, 21–29.

67. Lu, Z., Elliott, M.R., Chen, Y., Walsh, J.T., Klibanov, A.L., Ravichandran, K.S., and Kipnis, J. (2011). Phagocytic activity of neuronal progenitors regulates adult neurogenesis. Nat Cell Biol 13, 1076–1083.

68. Ma, S., Kwon, H.J., and Huang, Z. (2012). Ric-8a, a guanine nucleotide exchange factor for heterotrimeric G proteins, regulates bergmann glia-basement membrane adhesion during cerebellar foliation. The Journal of neuroscience : the official journal of the Society for Neuroscience 32, 14979–14993.

69. Ma, S., Santhosh, D., Kumar, T.P., and Huang, Z. (2017). A Brain-Region-Specific Neural Pathway Regulating Germinal Matrix Angiogenesis. Developmental cell 41, 366–381 e364.

70. Michaels, T.C.T., Saric, A., Curk, S., Bernfur, K., Arosio, P., Meisl, G., Dear, A.J., Cohen, S.I.A., Dobson, C.M., Vendruscolo, M., et al. (2020). Dynamics of oligomer populations formed during the aggregation of Alzheimer’s Abeta42 peptide. Nat Chem 12, 445–451.

71. Milosch, N., Tanriover, G., Kundu, A., Rami, A., Francois, J.C., Baumkotter, F., Weyer, S.W., Samanta, A., Jaschke, A., Brod, F., et al. (2014). Holo-APP and G-protein-mediated signaling are required for sAPPalpha-induced activation of the Akt survival pathway. Cell death & disease 5, e1391.

72. Moore, S.A., Saito, F., Chen, J., Michele, D.E., Henry, M.D., Messing, A., Cohn, R.D., Ross-Barta, S.E., Westra, S., Williamson, R.A., et al. (2002). Deletion of brain dystroglycan recapitulates aspects of congenital muscular dystrophy. Nature 418, 422–425.

73. Morley, J.E., Farr, S.A., Banks, W.A., Johnson, S.N., Yamada, K.A., and Xu, L. (2010). A physiological role for amyloid-beta protein:enhancement of learning and memory. J Alzheimers Dis 19, 441–449.

74. Muehlhauser, F., Liebl, U., Kuehl, S., Walter, S., Bertsch, T., and Fassbender, K. (2001). Aggregation-Dependent interaction of the Alzheimer’s beta-amyloid and microglia. Clin Chem Lab Med 39, 313–316.

75. Murase, S., Lantz, C.L., and Quinlan, E.M. (2017). Light reintroduction after dark exposure reactivates plasticity in adults via perisynaptic activation of MMP-9. eLife 6.

76. Murase, S., Winkowski, D., Liu, J., Kanold, P.O., and Quinlan, E.M. (2019). Homeostatic regulation of perisynaptic matrix metalloproteinase 9 (MMP9) activity in the amblyopic visual cortex. eLife 8.

77. Niewmierzycka, A., Mills, J., St-Arnaud, R., Dedhar, S., and Reichardt, L.F. (2005). Integrin-linked kinase deletion from mouse cortex results in cortical lamination defects resembling cobblestone lissencephaly. The Journal of neuroscience : the official journal of the Society for Neuroscience 25, 7022–7031.

78. Nishimoto, I., Okamoto, T., Matsuura, Y., Takahashi, S., Okamoto, T., Murayama, Y., and Ogata, E. (1993). Alzheimer amyloid protein precursor complexes with brain GTP-binding protein G(o). Nature 362, 75–79.

79. Pagenstecher, A., Stalder, A.K., Kincaid, C.L., Shapiro, S.D., and Campbell, I.L. (1998). Differential expression of matrix metalloproteinase and tissue inhibitor of matrix metalloproteinase genes in the mouse central nervous system in normal and inflammatory states. Am J Pathol 152, 729–741.

80. Palmeri, A., Ricciarelli, R., Gulisano, W., Rivera, D., Rebosio, C., Calcagno, E., Tropea, M.R., Conti, S., Das, U., Roy, S., et al. (2017). Amyloid-beta Peptide Is Needed for cGMP-Induced Long-Term Potentiation and Memory. The Journal of neuroscience : the official journal of the Society for Neuroscience 37, 6926–6937.

81. Pan, M., Xu, X., Chen, Y., and Jin, T. (2016). Identification of a Chemoattractant G-Protein-Coupled Receptor for Folic Acid that Controls Both Chemotaxis and Phagocytosis. Developmental cell 36, 428–439.

82. Papasergi-Scott, M.M., Stoveken, H.M., MacConnachie, L., Chan, P.Y., Gabay, M., Wong, D., Freeman, R.S., Beg, A.A., and Tall, G.G. (2018). Dual phosphorylation of Ric-8A enhances its ability to mediate G protein alpha subunit folding and to stimulate guanine nucleotide exchange. Science signaling 11.

83. Parodi, J., Sepulveda, F.J., Roa, J., Opazo, C., Inestrosa, N.C., and Aguayo, L.G. (2010). Beta-amyloid causes depletion of synaptic vesicles leading to neurotransmission failure. The Journal of biological chemistry 285, 2506–2514.

84. Patani, R., Hardingham, G.E., and Liddelow, S.A. (2023). Functional roles of reactive astrocytes in neuroinflammation and neurodegeneration. Nat Rev Neurol 19, 395–409.

85. Plant, L.D., Boyle, J.P., Smith, I.F., Peers, C., and Pearson, H.A. (2003). The production of amyloid beta peptide is a critical requirement for the viability of central neurons. The Journal of neuroscience : the official journal of the Society for Neuroscience 23, 5531–5535.

86. Preissler, J., Grosche, A., Lede, V., Le Duc, D., Krugel, K., Matyash, V., Szulzewsky, F., Kallendrusch, S., Immig, K., Kettenmann, H., et al. (2015). Altered microglial phagocytosis in GPR34-deficient mice. Glia 63, 206–215.

87. Puzzo, D., Privitera, L., Leznik, E., Fa, M., Staniszewski, A., Palmeri, A., and Arancio, O. (2008). Picomolar amyloid-beta positively modulates synaptic plasticity and memory in hippocampus. The Journal of neuroscience : the official journal of the Society for Neuroscience 28, 14537–14545.

88. Qin, H., Yeh, W.I., De Sarno, P., Holdbrooks, A.T., Liu, Y., Muldowney, M.T., Reynolds, S.L., Yanagisawa, L.L., Fox, T.H., 3rd, Park, K., et al. (2012). Signal transducer and activator of transcription-3/suppressor of cytokine signaling-3 (STAT3/SOCS3) axis in myeloid cells regulates neuroinflammation. Proceedings of the National Academy of Sciences of the United States of America 109, 5004–5009.

89. Ramaker, J.M., Swanson, T.L., and Copenhaver, P.F. (2013). Amyloid precursor proteins interact with the heterotrimeric G protein Go in the control of neuronal migration. The Journal of neuroscience : the official journal of the Society for Neuroscience 33, 10165–10181.

90. Ramsden, M., Henderson, Z., and Pearson, H.A. (2002). Modulation of Ca2+ channel currents in primary cultures of rat cortical neurones by amyloid beta protein (1-40) is dependent on solubility status. Brain research 956, 254–261.

91. Rice, H.C., Townsend, M., Bai, J., Suth, S., Cavanaugh, W., Selkoe, D.J., and Young-Pearse, T.L. (2012). Pancortins interact with amyloid precursor protein and modulate cortical cell migration. Development 139, 3986–3996.

92. Satz, J.S., Ostendorf, A.P., Hou, S., Turner, A., Kusano, H., Lee, J.C., Turk, R., Nguyen, H., Ross-Barta, S.E., Westra, S., et al. (2010). Distinct functions of glial and neuronal dystroglycan in the developing and adult mouse brain. J Neurosci 30, 14560–14572.

93. Sayed, F.A., Telpoukhovskaia, M., Kodama, L., Li, Y., Zhou, Y., Le, D., Hauduc, A., Ludwig, C., Gao, F., Clelland, C., et al. (2018). Differential effects of partial and complete loss of TREM2 on microglial injury response and tauopathy. Proceedings of the National Academy of Sciences of the United States of America 115, 10172–10177.

94. Schafer, D.P., and Stevens, B. (2015). Microglia Function in Central Nervous System Development and Plasticity. Cold Spring Harb Perspect Biol 7, a020545.

95. Shaked, G.M., Kummer, M.P., Lu, D.C., Galvan, V., Bredesen, D.E., and Koo, E.H. (2006). Abeta induces cell death by direct interaction with its cognate extracellular domain on APP (APP 597-624). FASEB J 20, 1254–1256.

96. Shankar, G.M., Li, S., Mehta, T.H., Garcia-Munoz, A., Shepardson, N.E., Smith, I., Brett, F.M., Farrell, M.A., Rowan, M.J., Lemere, C.A., et al. (2008). Amyloid-beta protein dimers isolated directly from Alzheimer’s brains impair synaptic plasticity and memory. Nat Med 14, 837–842.

97. Shigemoto-Mogami, Y., Hoshikawa, K., Goldman, J.E., Sekino, Y., and Sato, K. (2014). Microglia enhance neurogenesis and oligodendrogenesis in the early postnatal subventricular zone. The Journal of neuroscience : the official journal of the Society for Neuroscience 34, 2231–2243.

98. Spiller, K.J., Khan, T., Dominique, M.A., Restrepo, C.R., Cotton-Samuel, D., Levitan, M., Jafar-Nejad, P., Zhang, B., Soriano, A., Rigo, F., et al. (2019). Reduction of matrix metalloproteinase 9 (MMP-9) protects motor neurons from TDP-43-triggered death in rNLS8 mice. Neurobiol Dis 124, 133–140.

99. Spolidoro, M., Putignano, E., Munafo, C., Maffei, L., and Pizzorusso, T. (2012). Inhibition of matrix metalloproteinases prevents the potentiation of nondeprived-eye responses after monocular deprivation in juvenile rats. Cereb Cortex 22, 725–734.

100. Squarzoni, P., Oller, G., Hoeffel, G., Pont-Lezica, L., Rostaing, P., Low, D., Bessis, A., Ginhoux, F., and Garel, S. (2014). Microglia modulate wiring of the embryonic forebrain. Cell reports 8, 1271–1279.

101. Stellwagen, D., and Malenka, R.C. (2006). Synaptic scaling mediated by glial TNF-alpha. Nature 440, 1054–1059.

102. Stenman, J., Toresson, H., and Campbell, K. (2003). Identification of two distinct progenitor populations in the lateral ganglionic eminence: implications for striatal and olfactory bulb neurogenesis. The Journal of neuroscience : the official journal of the Society for Neuroscience 23, 167–174.

103. Stine, W.B., Jungbauer, L., Yu, C., and LaDu, M.J. (2011). Preparing synthetic Abeta in different aggregation states. Methods Mol Biol 670, 13–32.

104. Sturchio, A., Dwivedi, A.K., Malm, T., Wood, M.J.A., Cilia, R., Sharma, J.S., Hill, E.J., Schneider, L.S., Graff-Radford, N.R., Mori, H., et al. (2022). High Soluble Amyloid-beta42 Predicts Normal Cognition in Amyloid-Positive Individuals with Alzheimer’s Disease-Causing Mutations. J Alzheimers Dis.

105. Sturchio, A., Dwivedi, A.K., Young, C.B., Malm, T., Marsili, L., Sharma, J.S., Mahajan, A., Hill, E.J., Andaloussi, S.E., Poston, K.L., et al. (2021). High cerebrospinal amyloid-beta 42 is associated with normal cognition in individuals with brain amyloidosis. EClinicalMedicine 38, 100988.

106. Takamori, Y., Mori, T., Wakabayashi, T., Nagasaka, Y., Matsuzaki, T., and Yamada, H. (2009). Nestin-positive microglia in adult rat cerebral cortex. Brain research 1270, 10–18.

107. Tall, G.G., Krumins, A.M., and Gilman, A.G. (2003). Mammalian Ric-8A (synembryn) is a heterotrimeric Galpha protein guanine nucleotide exchange factor. The Journal of biological chemistry 278, 8356–8362.

108. Tan, J., Town, T., Paris, D., Mori, T., Suo, Z., Crawford, F., Mattson, M.P., Flavell, R.A., and Mullan, M. (1999). Microglial activation resulting from CD40-CD40L interaction after beta-amyloid stimulation. Science 286, 2352–2355.

109. Tran, N.M., Shekhar, K., Whitney, I.E., Jacobi, A., Benhar, I., Hong, G., Yan, W., Adiconis, X., Arnold, M.E., Lee, J.M., et al. (2019). Single-Cell Profiles of Retinal Ganglion Cells Differing in Resilience to Injury Reveal Neuroprotective Genes. Neuron 104, 1039–1055 e1012.

110. Tsiknia, A.A., Sundermann, E.E., Reas, E.T., Edland, S.D., Brewer, J.B., Galasko, D., Banks, S.J., and Alzheimer’s Disease Neuroimaging, I. (2022). Sex differences in Alzheimer’s disease: plasma MMP-9 and markers of disease severity. Alzheimers Res Ther 14, 160.

111. Vainchtein, I.D., Chin, G., Cho, F.S., Kelley, K.W., Miller, J.G., Chien, E.C., Liddelow, S.A., Nguyen, P.T., Nakao-Inoue, H., Dorman, L.C., et al. (2018). Astrocyte-derived interleukin-33 promotes microglial synapse engulfment and neural circuit development. Science 359, 1269–1273.

112. Walsh, D.M., Klyubin, I., Fadeeva, J.V., Cullen, W.K., Anwyl, R., Wolfe, M.S., Rowan, M.J., and Selkoe, D.J. (2002). Naturally secreted oligomers of amyloid beta protein potently inhibit hippocampal long-term potentiation in vivo. Nature 416, 535–539.

113. Wang, H., Ng, K.H., Qian, H., Siderovski, D.P., Chia, W., and Yu, F. (2005). Ric-8 controls Drosophila neural progenitor asymmetric division by regulating heterotrimeric G proteins. Nat Cell Biol 7, 1091–1098.

114. Wang, X., Jung, J., Asahi, M., Chwang, W., Russo, L., Moskowitz, M.A., Dixon, C.E., Fini, M.E., and Lo, E.H. (2000). Effects of matrix metalloproteinase-9 gene knock-out on morphological and motor outcomes after traumatic brain injury. The Journal of neuroscience : the official journal of the Society for Neuroscience 20, 7037–7042.

115. Wang, Y., Fu, W.Y., Cheung, K., Hung, K.W., Chen, C., Geng, H., Yung, W.H., Qu, J.Y., Fu, A.K.Y., and Ip, N.Y. (2021). Astrocyte-secreted IL-33 mediates homeostatic synaptic plasticity in the adult hippocampus. Proceedings of the National Academy of Sciences of the United States of America 118.

116. Yang, Y., Kim, J., Kim, H.Y., Ryoo, N., Lee, S., Kim, Y., Rhim, H., and Shin, Y.K. (2015). Amyloid-beta Oligomers May Impair SNARE-Mediated Exocytosis by Direct Binding to Syntaxin 1a. Cell reports 12, 1244–1251.

117. Yona, S., Kim, K.W., Wolf, Y., Mildner, A., Varol, D., Breker, M., Strauss-Ayali, D., Viukov, S., Guilliams, M., Misharin, A., et al. (2013). Fate mapping reveals origins and dynamics of monocytes and tissue macrophages under homeostasis. Immunity 38, 79–91.

118. Yoshida, M., Assimacopoulos, S., Jones, K.R., and Grove, E.A. (2006). Massive loss of Cajal-Retzius cells does not disrupt neocortical layer order. Development 133, 537–545.

119. Young-Pearse, T.L., Bai, J., Chang, R., Zheng, J.B., LoTurco, J.J., and Selkoe, D.J. (2007). A critical function for beta-amyloid precursor protein in neuronal migration revealed by in utero RNA interference. The Journal of neuroscience : the official journal of the Society for Neuroscience 27, 14459–14469.

120. Yu, D., Li, T., Delpech, J.C., Zhu, B., Kishore, P., Koshi, T., Luo, R., Pratt, K.J.B., Popova, G., Nowakowski, T.J., et al. (2022). Microglial GPR56 is the molecular target of maternal immune activation-induced parvalbumin-positive interneuron deficits. Sci Adv 8, eabm2545.

121. Zhang, X., Wang, Y., Supekar, S., Cao, X., Zhou, J., Dang, J., Chen, S., Jenkins, L., Marsango, S., Li, X., et al. (2023). Pro-phagocytic function and structural basis of GPR84 signaling. Nat Commun 14, 5706.

122. Zhang, Y., Chen, K., Sloan, S.A., Bennett, M.L., Scholze, A.R., O’Keeffe, S., Phatnani, H.P., Guarnieri, P., Caneda, C., Ruderisch, N., et al. (2014). An RNA-sequencing transcriptome and splicing database of glia, neurons, and vascular cells of the cerebral cortex. The Journal of neuroscience : the official journal of the Society for Neuroscience 34, 11929–11947.

123. Zhou, B., Lu, J.G., Siddu, A., Wernig, M., and Sudhof, T.C. (2022). Synaptogenic effect of APP-Swedish mutation in familial Alzheimer’s disease. Sci Transl Med 14, eabn9380.

124. Zipp, F., Bittner, S., and Schafer, D.P. (2023). Cytokines as emerging regulators of central nervous system synapses. Immunity 56, 914–925.

125. Zott, B., Simon, M.M., Hong, W., Unger, F., Chen-Engerer, H.J., Frosch, M.P., Sakmann, B., Walsh, D.M., and Konnerth, A. (2019). A vicious cycle of beta amyloid-dependent neuronal hyperactivation. Science 365, 559–565.

